# Rapid colorimetric detection of *citrus tristeza virus* combining portable sample preparation and reverse transcription-loop mediated isothermal amplification

**DOI:** 10.1101/2024.11.09.622765

**Authors:** Chia-Wei Liu, Sohrab Bodaghi, Manjunath L. Keremane, Brent Kalish, Georgios Vidalakis, Hideaki Tsutsui

## Abstract

A sensing platform combining semi-automated sample preparation protocol and one-step reverse transcription loop-mediated isothermal amplification (RT-LAMP) is reported for rapid colorimetric detection of *citrus tristeza virus* (CTV) in a greenhouse. An OmniLyse micro-homogenizer and cellulose paper disks were integrated for quick sample preparation of total nucleic acids (<15 min). RT-LAMP assays were optimized in terms of primers’ concentrations and minimization of false positives for both CTV and cytochrome oxidase (COX) detections. Specifically, the optimal reaction time for lab-based RT-LAMP assays was determined as 40 minutes with the detection limits of CTV and COX as 43 copies/μL (equivalent to 86 copies/mg of tissue) and 5 copies/μL (equivalent to 10 copies/mg of tissue), respectively. Additionally, an in-greenhouse colorimetric RT-LAMP assay with lyophilized reaction mix for endpoint CTV detection was successfully conducted in 35 minutes without a false response in either colorimetric or fluorometric assays. Overall, this quick sample preparation protocol integrated with the lyophilized RT-LAMP assays showed high efficiency and reliability in plant pathogen detection in a greenhouse. This strategy holds great potential to be integrated into a portable, autonomous system and be universally adopted for in-field diagnosis of different pathogens.

## 1. Introduction

Plant epidemics have long impacted crop yields and the global economy adversely, resulting in hundreds of billions of USD in crop yield losses every year [1-3]. The resulting food shortages further cause food crises across 59 countries/territories, affecting more than 280 million populations, as reported in the UN’s 2024 Global Report on Food Crises [4]. Additionally, worsening climate conditions such as frequent high temperatures and drought will further exacerbate the current situation[5]. As an example, citrus, one of the most popular cash crops in the world, was once affected by *citrus tristeza virus* (CTV). CTV, a phloem-limited RNA virus, causes a number of devastating diseases, such as yellow dwarf and lime dieback, killing more than 70 million citrus plants worldwide in the past few decades [6-9]. CTV has multiple symptoms, including quick decline, seedling yellows, growth cutoff, stunting, and stem pitting with low yield and poor-quality citrus fruits [10]. To alleviate this situation, researchers have devised many strategies to combat CTV, including selective eradication of infected plants identified based on serological and genotypic assays, reproduction of virus-free plants, and controlling viruliferous vectors using pesticides [11, 12]. In addition, grafting desired scions onto sour orange rootstocks, tolerant against infections, is another common strategy [13]. However, these strategies cannot be effectively utilized if diseased trees cannot be accurately identified from a pool of healthy ones promptly. Therefore, a platform allowing an efficient and accurate diagnosis in the field is critically important for better disease management.

The current protocols for plant disease diagnostics rely on specimen collection and submission by growers, who need to collect multiple specimens from suspected trees, place them into clean, individually labeled containers, and mail them to centralized labs for diagnosis [14]. Any damage due to temperature or distance during transit could result in unsuccessful identification (e.g., false negatives). At the lab, a series of specimen processing steps (manual tissue homogenization and nucleic acid extraction) and the subsequent molecular (e.g., PCR/qPCR) and genotypic (e.g., NGS) assays could take anywhere from a few days to weeks, depending on the specimen amounts [15]. Despite the high-quality results provided by these commercial tools, high start-up costs, and expensive materials could make routine testing unfeasible as demand increases [16-18].

Even field-ready serological tests such as lateral flow immunoassays (LFIAs) that are frequently used for prescreening of target plant diseases in the field (e.g., AgriStrip, Pocket Diagnostics kits, etc.) still require follow-up testing for samples tested positive with more specific techniques (e.g., qPCR, pathogen culture) in advanced labs [16, 19, 20]. Additionally, a standard way of manual sample homogenization using mortar and pestle requires continuous addition of liquid nitrogen while grinding and skillful personnel to avoid potential human errors. This is usually time-consuming and inaccessible outside labs. Although using high-throughput machines, such as BeadBlaster 24 Microtube Homogenizer, RETSCH MM 400 Mixer Mill, and Thermo Scientific KingFisher Flex Purification System, effectively save time and effort by automating large-scale homogenization and liquid handling steps, these equipment and kits are still unaffordable for low-infrastructure sites in developing countries, let alone in the field [21, 22]. Thus, an affordable platform allowing in-field deployment for rapid and sensitive detection is of great importance for better plant disease management.

To address the bottleneck and benefit in-field disease management, mechanical lysis using portable techniques, such as microneedle patch and 3D-printed handheld device, have been adopted for quick sample preparation [23, 24]. Specifically, Claremont BioSoultions’ OmniLyse micro-homogenizer has been used as an effective approach to not only produce plant lysates [25] but also quickly extract total nucleic acids with the use of cellulose paper [26]. Such mechanical lysis, combined with equipment-free nucleic acid extraction (e.g., cellulose paper), has been proven sufficient for total nucleic acid extraction directly from plant lysates and is suitable for in-field use due to their convenient-to-use features and manageability [27-30]. To increase the reliability of disease diagnosis in the field, loop-mediated isothermal amplification (LAMP) has emerged as a sensitive and reliable diagnostic tool that does not require complicated thermo-cycling (isothermal amplification at 63–65°C), has a short reaction time (30-40 minutes), and has a high tolerance to impurities [31, 32]. So far, scientists have developed an integrated diagnosis platform combining a microneedle patch for quick nucleic acid extraction, on-chip LAMP/reverse transcription (RT)-LAMP assays, and smartphone-based image analysis for the detection of multiple tomato pathogens [33, 34]. Despite the short turnaround time and high sensitivity, that platform has only been verified with tomatoes, as the microneedle patch is most suitable for tomato’s thin and soft leaves. However, LAMP itself was proven to be a reliable alternative to qPCR-based assay for in-field plant disease diagnosis.

As a follow-up improvement of our previous study [26], we present a rapid and sensitive platform for the visual detection of CTV by combining a micro-homogenizer, chromatography paper, and RT-LAMP colorimetric assay (Figure 1). We adapted our previously developed protocol for quick sample preparation [26], followed by RT-LAMP assays, in which the primer design, primer concentrations, and reaction times were optimized. Additionally, the role of Dimethyl sulfoxide (DMSO) in minimizing false positives was investigated, and optimal concentrations were determined. To further enable in-field plant disease diagnosis, lyophilization of the RT-LAMP reaction mix was attempted and compared side by side with freshly prepared controls. Consequently, colorimetric RT-LAMP assays with lyophilized RT-LAMP reaction mix were performed with optimal conditions in a greenhouse for CTV diagnosis.

**Fig. 1.**
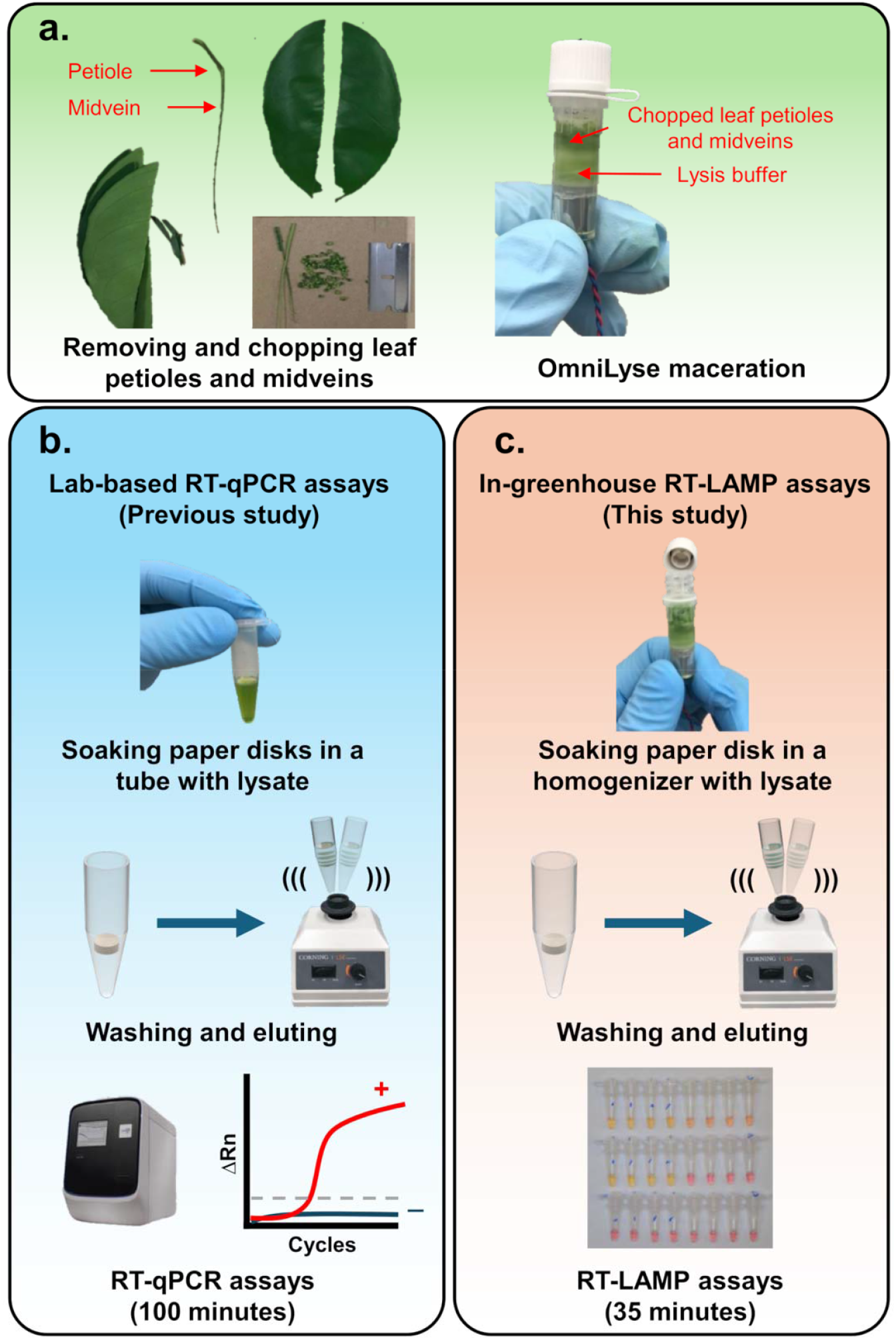
Stepwise workflow starting from (a) sample collection and processing to (b) lab-based nucleic acid extraction and RT-qPCR [26] and (c) in-field nucleic acid extraction and detection utilizing colorimetric RT-LAMP (present study).

## 2. Materials and methods

### 2.1 Materials and equipment

Guanidine isothiocyanate (GITC), polyvinylpyrrolidone (PVP), Dimethyl sulfoxide (DMSO), D-(+)-Trehalose dihydrate, guanidine hydrochloride (GuHCl) and Whatman Grade 1 CHR cellulose chromatography papers (Cat. No. 3001-861) were purchased from Sigma-Aldrich (St. Louis, MO). Sodium acetate trihydrate (pH 5.0), sodium chloride, ethylenediaminetetraacetic acid (EDTA), Tris-EDTA 1X solution (TE, pH 8.0), Tris-borate-EDTA (TBE) solution and MagMAX-96 Viral RNA isolation kit (Cat. No. AM1836) were purchased from Thermo Fisher Scientific (Waltham, MA). 1M Tris-HCl solution (pH 7.5) (Cat. No. 15567-027) was purchased from Invitrogen (Waltham, MA). AgPath-ID One-Step RT-PCR kit was purchased from Life Technologies (Carlsbad, CA). WarmStart Colorimetric LAMP kit (Cat. No. M1800S/L), gel loading dye (10X), and 100 bp DNA ladder were purchased from New England BioLabs (Ipswich, MA). Green Fluorescent Dye (Item ID 30078-1) was purchased from LGC Biosearch Technologies (Hoddesdon, UK). Primers and probes for RT-qPCR and RT-LAMP assays were synthesized by Integrated DNA Technologies (Coralville, IA). Nuclease-free water was used to prepare all buffers and MasterMix throughout the study.

Devices and lab equipment-wise, MagMAX Express-96 Magnetic Particle Processor, NanoDrop 2000c spectrophotometer, and Applied Biosystems QuantStudio 12K Flex Real-Time PCR System were purchased from Thermo Fisher Scientific (Waltham, MA). Allegra X-22R Centrifuge was purchased from Beckman Coulter (Indianapolis, IN). CFX Connect Real-Time PCR Detection System and PowerPac Basic Electrophoresis Power Supply and ChemiDoc Imaging System were purchased from BIO-RAD (Hercules, CA). FreeZone Triad Cascade Benchtop Freeze Dryer was purchased from Labconco (Kansas City, MO). OmniLyse X-mini Cell & Tissue Lysis Device (Item# 01.532.48), also referred to as OmniLyse micro-homogenizer) was purchased from Claremont BioSolutions (Upland, CA).

### 2.2 Preparation of buffers

Lysis buffer was prepared with 4 M GITC, 0.2 M sodium acetate trihydrate, 2 mM EDTA, and 2.5 % PVP, with the final pH adjusted to 5.0 using 1N HCl solution [26]. Wash buffer was prepared by diluting 1 M Tris-HCl solution to 10 mM for washing paper disks after soaking in the crude lysate. The elution buffer used for the benchtop extraction protocol was TE 1X solution, whereas an additional 0.1 M sodium chloride was added specifically for eluting nucleic acids from paper disks [26].

### 2.3 Plant materials and sample collection

Fourteen CTV-infected samples from different citrus varieties (Table S1) were collected to determine the specificity of the RT-LAMP primers. Healthy sour orange (*Citrus aurantium* L.) was used throughout the study as a negative control to evaluate the developed protocols. These sources were all provided by the Citrus Clonal Protection Program (CCPP) disease bank at the University of California, Riverside. All tested leaves were evenly collected on the east, west, south, and north sides of the tree at eye level to ensure consistency of sampling. Before processing, the leaves were gently wiped with wet paper towels to remove any dust.

#### 2.4 Benchtop protocol for sample preparation

### 2.4.1 Preparation of crude lysates

Phloem-rich petioles and midveins removed from freshly picked healthy and diseased leaves were ground using a mortar and pestle with a gradual addition of liquid nitrogen until finely pulverized. The tissues were then aliquoted to 250 mg each in 2 mL centrifuge tubes, lysed by mixing with 750 µL of the lysis buffer and incubating for 15 min at 4°C. The mixture was centrifuged at 14,000 × g at 4°C for 25 min, after which 150 μL of the supernatant was collected for subsequent nucleic acid extraction.

#### 2.4.2 Nucleic acid extraction using benchtop equipment

Nucleic acids in crude lysates were isolated using MagMAX Express-96 Magnetic Particle Processor (previously programmed) with a MagMAX-96 Viral RNA isolation kit. Each reaction contained 150 μL of supernatant (crude lysate), 22 μL of RNA binding bead mix (10 μL lysis/binding enhancer + 10 μL RNA binding beads + 2 μL carrier RNA), 139 μL of lysis/binding solution as recommended by the manufacturer and extra 139 μL of 100% isopropanol as previously reported by Dang et al. [15]. The resulting nucleic acids were eluted in 100 μL of TE 1X solution and were quickly assessed using the NanoDrop 2000c spectrophotometer to confirm the quality before being aliquoted and stored at −80°C.

### 2.5 Semi-automated protocol for sample preparation

#### 2.5.1 Preparation of crude lysates

As introduced and optimized in our previous work [26], OmniLyse X-mini Cell & Tissue Lysis Devices were used to macerate 100 mg of chopped petioles and midveins from freshly picked healthy and diseased leaves, along with 300 mg of ceramic microbeads and 300 μL of lysis buffer for 5-min lysis at 11 V. The resulting crude lysate (250 μL) was drawn out using a filtered syringe tip and collected in a centrifuge tube for subsequent nucleic acid extraction.

#### 2.5.2 Nucleic acids extraction using paper disks

As optimized in our previous work [26], five paper disks (d = 6.35 mm), made of chromatography paper, were subject to a 5-minute soaking in 250 µL of crude lysate for rapid nucleic acid capturing. Disks were then transferred to a clean centrifuge tube containing 1 mL of 10 mM Tris-HCl solution for a 1-minute wash with mild vortex. These washed disks were suitable for either immediate elution in 100 µL of modified TE 1X solution for molecular assays or storage at room temperature for later use after being fully air dried.

### 2.6 RT-qPCR assays

Primers and probes were adapted from previous reports, with their quality and homology validated [35]. Cytochrome oxidase (COX) genes in plant mitochondrial genomes are routinely used as a reliable internal control to confirm nucleic acid integrity. The sequences and the corresponding nucleotide positions of these primers and probes are listed in Table 1. All the assays were conducted in an Applied Biosystems QuantStudio 12K Flex Real-Time PCR System, as described earlier [26].

**Table 1.**
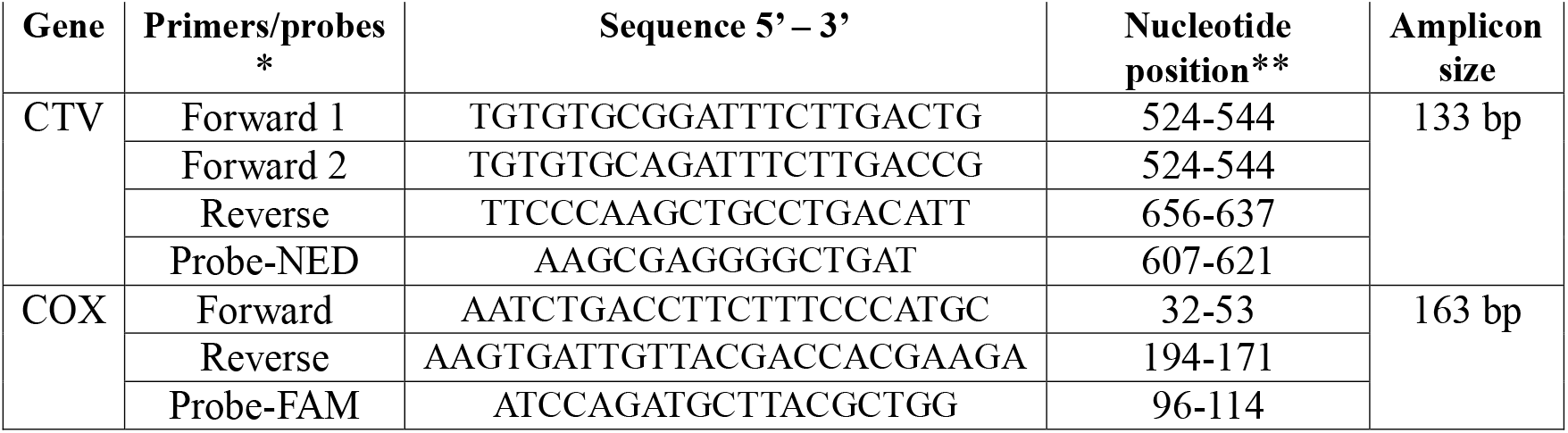
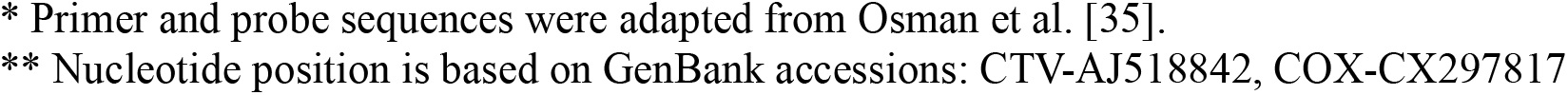
CTV and COX primers and probes for RT-qPCR assays.

### 2.7 RT-LAMP assays

Two primer sets, with six specific primers (F3/B3, FIP/BIP, and LB/LF) in each, were designed using PrimerExplorer v5 (Eiken Chemical, Japan) and subject to BLAST analysis to confirm their specificities. The CTV primers were designed based on the conserved region determined through proper sequence alignment (Table S1). A similar approach was used for designing COX primers (Table S2). The sequences of RT-LAMP primer sets are detailed in Table 2, while the overall comparison of nucleotide positions between RT-qPCR and RT-LAMP primers is available in Table. S1 and S2. The protocols for preparing the RT-LAMP reaction mixture were also modified to adapt to various needs, which are detailed below. All the assays were conducted isothermally _at_ 65°C either in a CFX Connect Real-Time PCR Detection System or a portable dry block heater to fit specific purposes.

**Table 2.**
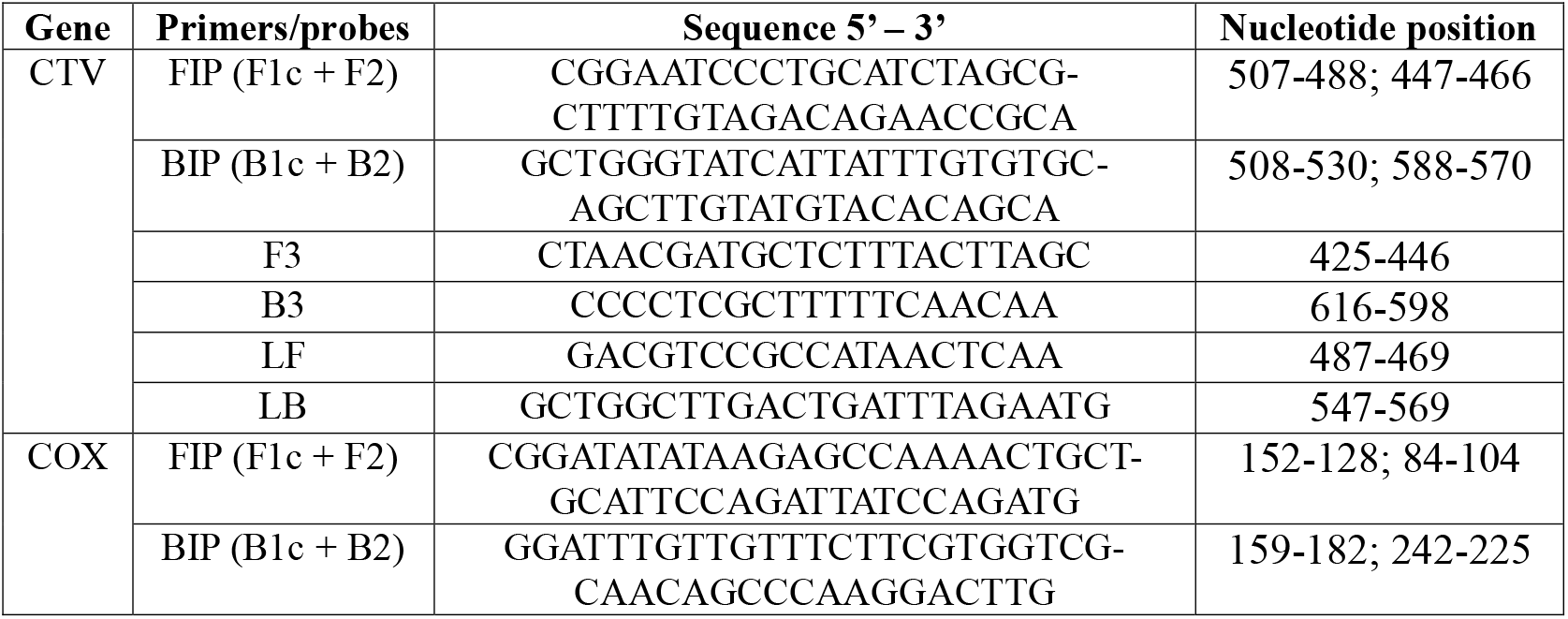
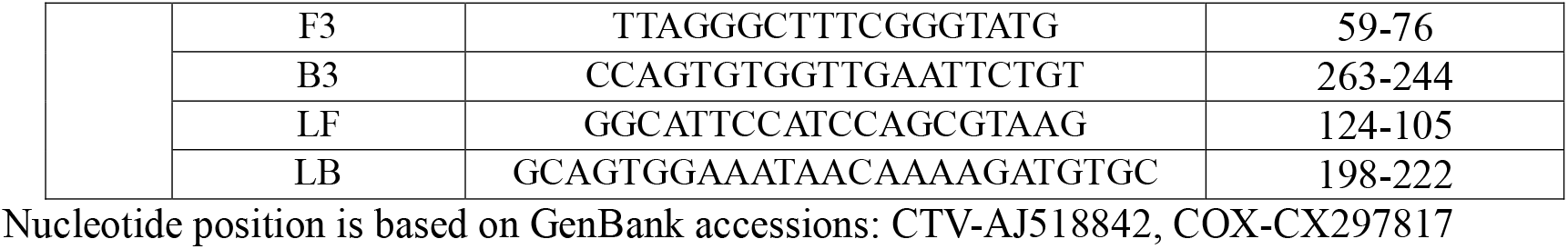
CTV and COX primers for RT-LAMP assays.

#### 2.7.1 Preparation of DMSO-based RT-LAMP reaction mixture

The original protocol suggested by the manufacturer was modified below [36]. Each reaction contained 12.5 μL of 2X Master Mix, 2.5 μL of 10X Primer Mix (final concentration: 1.6 μM of FIP/BIP, 0.2 μM of F3/B3 and 0.75 μM LF/LB), 1.5 μL of DMSO, 1.25 μL of Green Fluorescent Dye, 5.25 μL of nuclease-free water, and 2 μL of RNA template (or nuclease-free water for non-target control). Green Fluorescent Dye was added for visualization via CFX Connect Real-Time PCR Detection System.

#### 2.7.2 Preparation of lyophilized RT-LAMP reaction mixture

The lyophilization protocol previously developed by Song et al. [37] was adopted in the study. Each reaction contained 12.5 μL of 2X Master Mix, 2.5 μL of 10X Primer Mix (final concentration: 1.6 μM of FIP/BIP, 0.2 μM of F3/B3 and 0.75 μM LF/LB), 5.75 μL of D-(+)-Trehalose dihydrate solution (4.296 g/10 mL), 1.25 μL of Green Fluorescent Dye, 1 μL of GuHCl solution (0.955 g/10 mL) and 2 μL of RNA template (or nuclease-free water for non-target control). The reaction mixtures in 1.5 mL Eppendorf tubes were placed at −80°C until fully frozen before being freeze-dried for 18-20 hours in a FreeZone Triad Cascade Benchtop Freeze Dryer. The resulting freeze-dried powders were then stored either at −80°C or in a sealed Ziploc with desiccant at room temperature for later use.

### 2.8 Gel electrophoresis

Amplicons from RT-qPCR and RT-LAMP were visualized by agarose gel electrophoresis (2% w/v) in 0.5X TBE solution. Amplified samples were mixed with gel loading dye (10X) and loaded into the wells. A 100 bp DNA ladder was used as a size marker (M). The gel was run for 30 min at 100 V using a PowerPac Basic Electrophoresis Power Supply. The resultant gel was immersed in ethidium bromide solution (0.06% v/v) with gentle shaking for 10 min for DNA post-staining and was imaged using a ChemiDoc Imaging System.

## 3 Results and discussion

### 3.1 Characterization of RT-LAMP primers

#### 3.1.1 Optimization of primer concentrations for CTV RT-LAMP assays

To determine the desired reaction time for CTV RT-LAMP assays, CTV isolate T529 (accession no. KC841789.1) was selected as the target for this set of tests due to its 100% consistency to the designed primers. The RT-LAMP assays were carried out at 65°C with varying concentrations of inner primer (1.2 μM, 1.6 μM, and 2 μM), loop primer (0.5 μM, 0.75 μM, and 1 μM) and a fixed outer primer concentration (0.2 μM) in different combinations [38]. As presented in Table 3, all the combinations showed nearly equivalent threshold times, ranging from 9.6 to 10.4 minutes. Particularly, the combination with 1.6 μM of inner primers, 0.2 μM of outer primers, and 0.75 μM of loop primers (bolded in Table 3) performed almost one minute shorter in threshold time (09’33”) than the longest one (10’21”). Thus, this primer combination was chosen and used throughout the study for the preparation of both CTV and COX primer mixtures.

**Table 3.** Optimization of primer concentrations for RT-LAMP assay for CTV.

| Inner primers<br>(FIP/BIP) (µM) | Outer primers<br>(F3/B3) (µM) | Loop primers<br>(LF/LB) (µM) | Threshold time*<br>(min:sec) |  |  | Mean Time<br>(min:sec) | SD |
| --- | --- | --- | --- | --- | --- | --- | --- |
|  |  |  | R1 | R2 | R3 |  |  |
| 1.2 | **0.2** | 0.5 | 09:51 | 09:41 | 09:35 | 09:43 | 0:08 |
| 1.6 |  |  | 09:38 | 09:37 | 09:42 | 09:39 | 0:02 |
| 2 |  |  | 10:04 | 10:15 | 10:16 | 10:12 | 0:07 |
| 1.2 |  | **0.75** | 09:39 | 09:34 | 09:43 | 09:39 | 0:05 |
| **1.6** |  |  | **09:36** | **09:36** | **09:27** | **09:33** | **0:05** |
| 2 |  |  | 10:28 | 10:20 | 09:57 | 10:15 | 0:16 |
| 1.2 |  | 1 | 09:46 | 09:46 | 09:38 | 09:43 | 0:05 |
| 1.6 |  |  | 10:01 | 09:51 | 09:57 | 09:56 | 0:05 |
| 2 |  |  | 10:18 | 10:29 | 10:17 | 10:21 | 0:07 |

\*Threshold time (Tt) represents the time at which the fluorescence was equal to the threshold value during LAMP assays.

#### 3.1.2 Minimization of false positives for CTV RT-LAMP assays

As previously reported by Shao et al.,[39] DMSO has been frequently used to inhibit secondary structures in DNA templates and primers and also to interfere with self-complementarity of the DNA in PCR and LAMP assays, resulting in fewer false positives. Thus, RT-LAMP reaction mixtures with varying concentrations of DMSO (0%, 4%, 6%, and 8% (v/v)) added were tested separately with CTV positive controls (PC, N = 3), negative controls (N C, N = 8), and non-target controls (NTC, N = 8) and incubated at 65°C for 80 minutes. As depicted in Fig.2, the addition of DMSO significantly delayed the threshold times of all controls, including PC, by inhibiting the interactions between templates and primers, as well as polymerase activity [40]. To optimize the detection window for RT-LAMP assays, the time difference between the average threshold times of PCs and the earliest false positive was compared across different DMSO concentrations (0%, 4%, 6%, and 8%). The RT-LAMP assays with 6% DMSO had the broadest detection window, significantly delaying the occurrence of false positives compared to the assay with 0% and 4% DMSO. However, increasing the DMSO concentration to 8% did not statistically enhance the suppression of false positives compared to the 6%, while it extended the threshold time of true positives and shortened the detection window. This suggests diminishing returns with higher DMSO concentrations. Based on the results, the addition of 6% DMSO was selected as the optimal protocol for the preparation of RT-LAMP mixtures throughout the study. A 40-minute line was considered a proper reaction time for our lab protocols, within which the least false positives were observed.

**Fig. 2.**
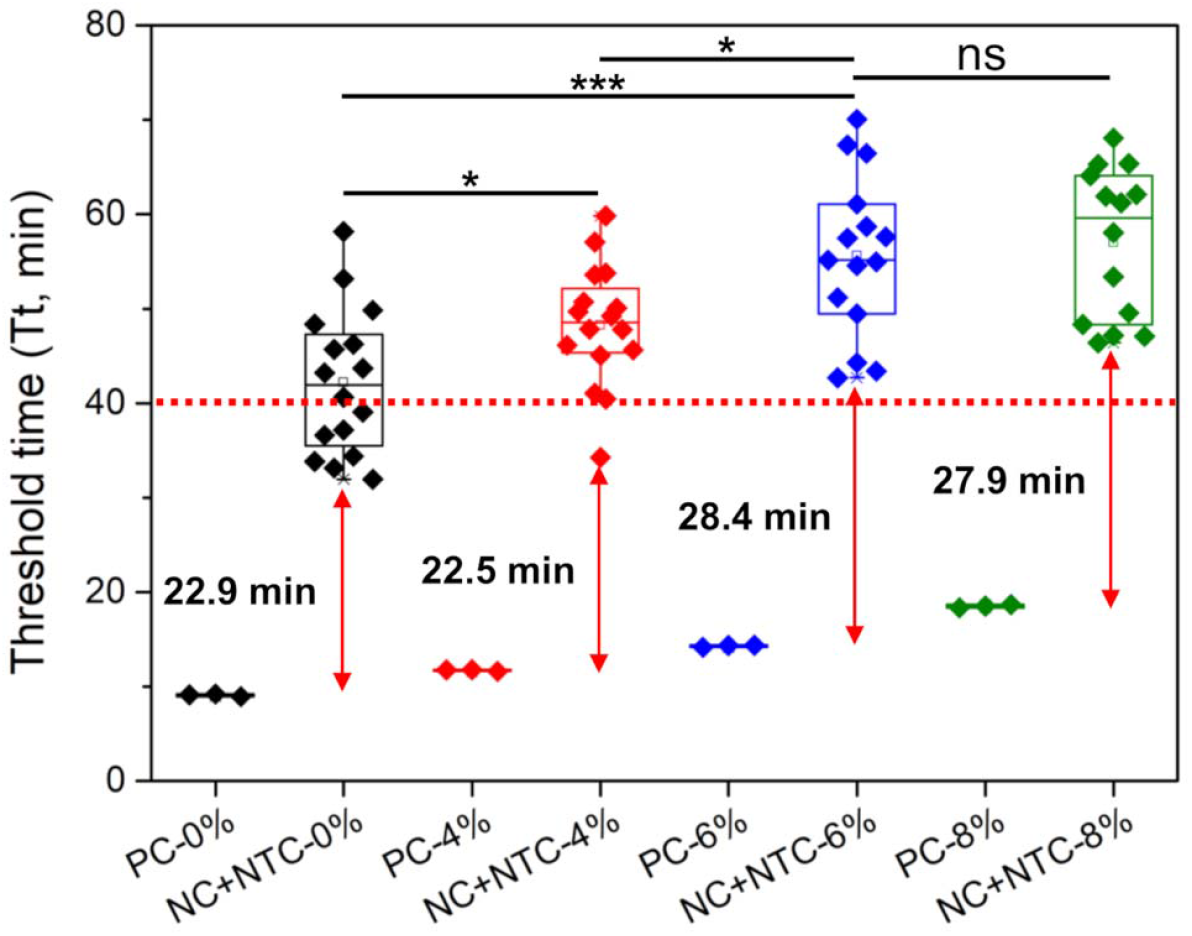
Threshold time comparison among RT-LAMP assays with 0%, 4%, 6%, and 8% DMSO (N = 3 for PC, N = 16 for NC and NTC combined). For example, PC-0% is CTV positive control with 0% DMSO, and NC+NTC-8% is the combination of negative control with 8% DMSO and no-target control with 8% DMSO. The 6% and 8% DMSO conditions did result in 1 and 2 true negatives, respectively, where no signal was present after the entire 80-minute reaction window. *and *** denote p ≤ 0.05 and p ≤ 0.001, respectively; ns (not significant) denotes p > 0.05.

#### 3.1.3 Evaluation of the detection limit for CTV and COX RT-LAMP assays

To evaluate the detection limit of the RT-LAMP primers for CTV and COX, total nucleic acids of a CTV-positive source (isolate T529) were selected as a positive control and serially diluted to 10^−6^ for RT-qPCR assays. As presented in Fig. 3a and 3c, the resulting formulas of the RT-qPCR standard curves for CTV and COX were produced and used to estimate the detection limits of RT-LAMP assays. These curves presented good linearity within the detection limit, revealing high correlations between Cq and cDNA quantities (R^2^ > 0.99 for both genes). RT-LAMP assays were then conducted with the same set of samples. It was observed that serial dilutions 10^−5^ and 10^−6^ showed false negatives in CTV RT-LAMP assay, whereas the detection limit of COX RT-LAMP assay was on the order of 10^−5^. The results showed that the RT-LAMP assays were able to achieve detection limits of 43 copies/μL (equivalent to 86 copies/mg of tissue) and 5 copies/μL (equivalent to 10 copies/mg of tissue) for CTV and COX, respectively (Fig. 3b and 3d). The calculation process is detailed in Table. S3. Standard curves of RT-LAMP assays were also generated with high linearity over at least five orders of magnitude (R^2^ = 0.95 for CTV and R^2^ = 0.98 for COX). Although the detection limit of RT-LAMP assay was approximately 1-2 orders of magnitude higher than that of RT-qPCR, target RNA with 10^5^ – 10^6^ copies/2 μL can be detected within a short period of time (16–17 minutes). It is also expected that, within the detection range, the most diluted samples can be detected within 28 minutes based on the derived formulas (when Log10(copies) = 0). Additionally, the gel patterns of both the RT-qPCR and RT-LAMP assays indicate correct amplification, as does the color change of the RT-LAMP tubes (Fig. S1).

**Fig. 3.**
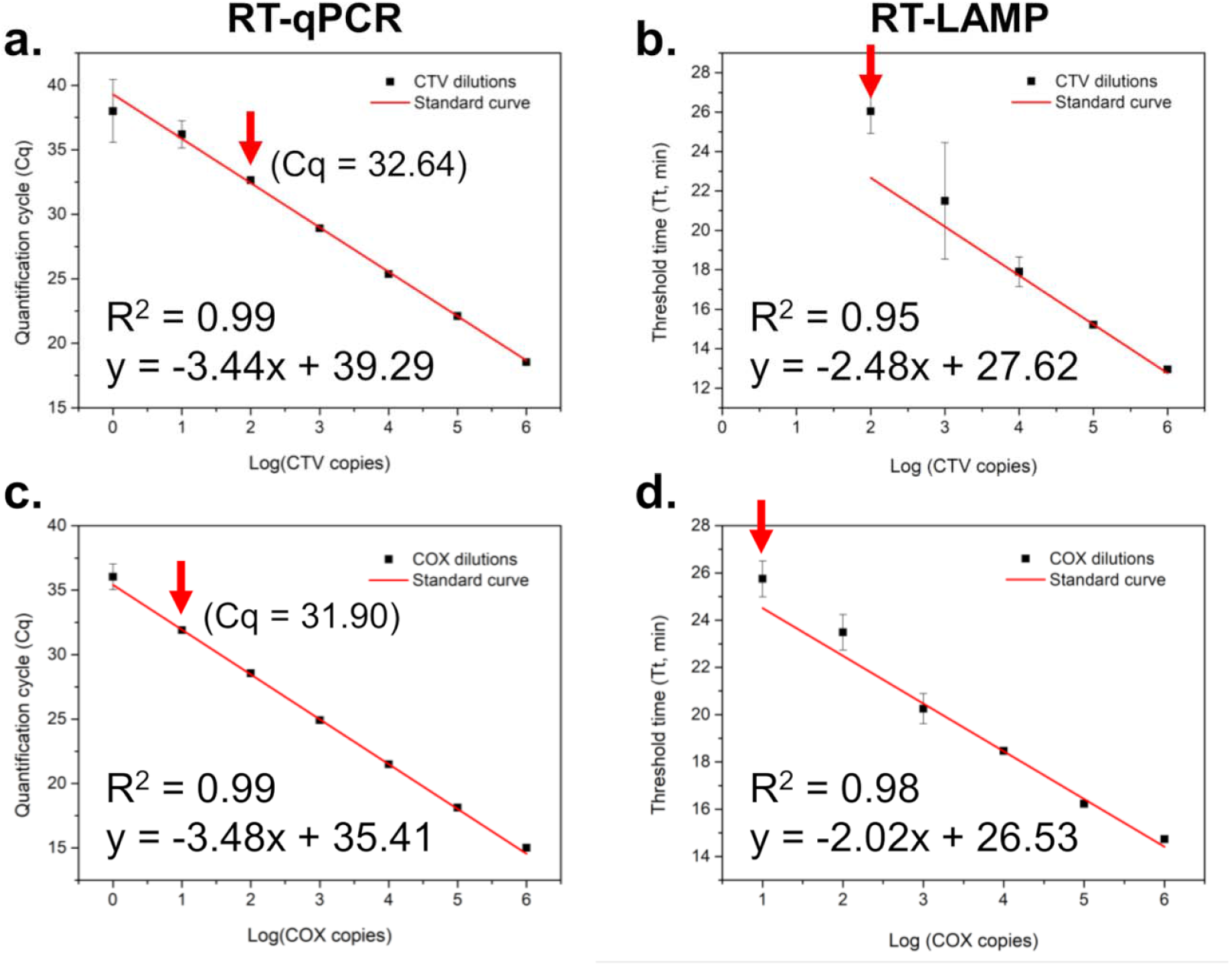
Sensitivity comparison of RT-qPCR and RT-LAMP assays using 10-fold serially diluted total nucleic acids of a CTV-positive source (T529), containing 10^0^ – 10^−6^ dilutions. (a) Standard curve derived from CTV RT-qPCR assays (N = 3), (b) Standard curves derived from CTV RT-LAMP assays (N = 3), (c) Standard curve derived from COX RT-qPCR assays (N = 3), and (d) Standard curve derived from COX RT-LAMP assays (N = 3). The red arrowhead indicates the detection limits of RT-LAMP assays for CTV and COX, and its corresponding dilution in the RT-qPCR assays.

### 3.2 Assessment of the RT-LAMP detection efficacy to various CTV isolates

Specifically, plants with CTV isolates aligned with the designed primers, as listed in Table S4, were collected for the test from the CCPP greenhouses at the University of California, Riverside. To determine the presence of viruses, total nucleic acids were extracted from these sources according to the standard lab protocol (section 2.4), and RT-qPCR was performed in triplicates (Table S4). Similarly, the RT-LAMP reactions of the same sources were also carried out and compared side by side with those of RT-qPCR (Table S4). The results demonstrated that the RT-LAMP CTV reactions functioned with equal efficiency as the standard lab protocol for the selected isolates. Additionally, the COX genes in these isolates were all detected by the RT-LAMP COX primers in the assays (Table S4), proving good sensitivity and specificity. Although the CTV primer set is not broadly applicable to all the isolates, it was still sufficient for our demonstration and improvements of the RT-LAMP protocol and its implementation in the field.

### 3.3 In-greenhouse RT-LAMP assays

#### 3.3.1 Evaluation of lyophilized RT-LAMP reagents

To further adapt the current RT-LAMP protocol for in-field application, it is critically important to streamline the procedures by having all the required reagents, except for the viral template, mixed in a tube in advance. Furthermore, reagents that can be stored and shipped under less stringent conditions, such as at room temperature, are even more ideal, particularly for diagnostic purposes in resource-limited regions. Given these requirements, lyophilization of the RT-LAMP reagents was considered an appealing solution. It has been reported that trehalose, a disaccharide, provides cryo- and lyoprotection for proteins (e.g., polymerase) during the lyophilization process and enhances stability during long-term storage [41]. Further, the addition of guanidine hydrochloride in the mixture helps expedite the reaction and improve the sensitivity of colorimetric RT-LAMP assays [42]. Therefore, a previously reported protocol by Song et al. [37] was adapted for the preparation of lyophilized RT-LAMP reagents. To evaluate the performance of the lyophilized reaction mix after one-week storage, the resulting lyophilized mixtures were stored at −80°C and room temperature separately. Serving as controls for comparison, the original DMSO-based mixture and fresh trehalose-based control were also included to evaluate the threshold time of PC and the false positive rate within a 40-minute window. The images depicted the lyophilized reaction mix before and after one-week storage, and the comparison between fresh controls and rehydrated reaction mix are all detailed in Fig. S2.

As depicted in Fig. 4a, trehalose-based assays for CTV generally resulted in earlier PC thresholds and no significant difference in NCs+NTCs thresholds compared to DMSO-based assays, although a false positive in lyophilized reaction mix (−80°C) and in fresh trehalose control were found within the highlighted 35-minute region and 40-minute line. Fewer false positives, as annotated in brackets (true negative rate), were also found in these trehalose-based assays, indicating the combination of trehalose and guanidine hydrochloride is a promising alternative to DMSO without a negative impact on PC’s threshold time. In RT-LAMP assays of COX (Fig. 4b), a false positive in the lyophilized reaction mix (room temp) was found very close to its PC’s threshold, while the rest showed no significant difference between each other. Similarly, true negatives were also found in most of these reactions, including DMSO-based assays. Notably, regardless of storage conditions, trehalose-based assays with a lyophilized reaction mix showed no significant differences from the fresh control for CTV and COX assays. Therefore, room temperature storage was selected for our next study. Although trehalose-based assays overall demonstrated better outcomes with fewer false positives and earlier PC thresholds, random false positives within the 35 and 40-minute window (as highlighted in Fig. 4a, b) could still be an uncertainty for in-field application, leading to unreliable detection.

**Fig. 4.**
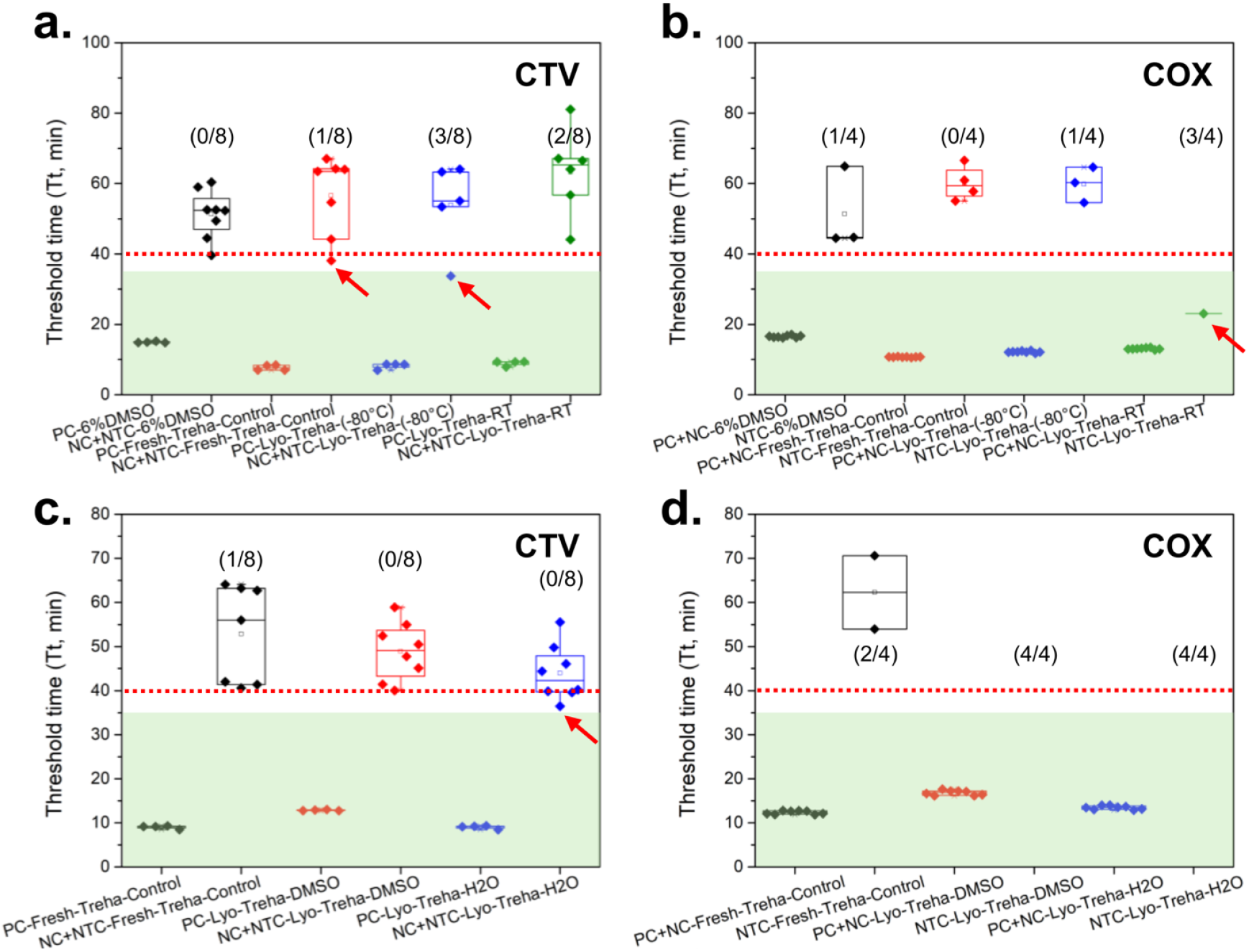
Evaluation of RT-LAMP assays with lyophilized reaction mix (N = 4). Comparison of threshold times between (a) fresh controls and H2O-rehydrated reaction mix in RT-LAMP CTV assays, (b) fresh controls and H2O-rehydrated reaction mix in RT-LAMP COX assays, (c) trehalose fresh control and H2O-rehydrated and DMSO-rehydrated reaction mix in RT-LAMP CTV assays, and (d) trehalose fresh control and H2O-rehydrated and DMSO-rehydrated reaction mix in RT-LAMP COX assays. The red dotted line represents the 40-minute window, whereas the green highlighted region refers to the 35-minute window.

To eliminate false positives within the reaction window, a 6%-DMSO solution was used to rehydrate the lyophilized reaction mix stored at room temperature and was compared with the fresh trehalose-based control and the H2O-rehydrated reaction mix. As presented in RT-LAMP CTV assays (Fig. 4c), the lyophilized mix rehydrated with 6%-DMSO solution delayed false negatives for 3.5 minutes, whereas the first false positives in DMSO-rehydrated assays and the fresh control almost happened at the same time. Notably, in COX assays (Fig. 4d), no false positives were found in either rehydrated assay, while 2 out of 4 were still found in the fresh control. This set of tests suggested that the addition of DMSO effectively suppressed false positives from happening within the 40-minute window. Although H2O-rehydrated assays also showed no false positives in the test, the addition of DMSO is still recommended for a more reliable outcome.

Overall, RT-LAMP assays with H2O-rehydrated lyophilized reaction mix showed no difference in PC threshold times, regardless of storage conditions. Rehydration using a 6% DMSO solution further delayed the false positive threshold, ensuring reliable detection within the designated reaction time. Given the reported outcomes, a lyophilized reaction mix (room temperature) rehydrated with 6%-DMSO solution (referred to as modified trehalose-based protocol) was then selected for in-greenhouse CTV RT-LAMP assays in the next section.

#### 3.3.2 Implementation of in-greenhouse colorimetric RT-LAMP assays

Since the modified trehalose-based protocol was determined to be as reliable as the lab-based assays, our next attempt was to conduct onsite CTV RT-LAMP assays in a greenhouse using the previously reported micro-homogenizer and cellulose paper protocols. Leaves were directly collected from a CTV-positive plant (isolate T520) and a CTV-negative sweet orange for onsite sample processing. The lyophilized reaction mix for CTV and COX detection were rehydrated on the spot and temporarily placed on ice before use. Total nucleic acids were eluted from soaked paper disks after washing and were also placed on ice for later use. RT-LAMP assays were conducted by inserting the tube strips in a heat block for 28 and 35 minutes, respectively, to avoid any possible false positives and shorten the reaction time. The tube strips were then placed on ice immediately after incubation to halt the reaction.

As depicted in Fig. 5a left, the CTV-positive, CTV-negative, and water control were ordered sequentially from top to bottom for both CTV and COX RT-LAMP assays (N = 4). The color changes were distinguishable for all the tests after 28-minute incubation at 65°C, although the color in the positive reactions remained slightly orangish, implying the reaction was not fully completed. In addition, differences in fluorescence readouts between positive and negative reactions (Fig. 5a right) were consistent, although the change was not as obvious as seen in the colorimetric assays. Further, in the longer incubation (35 minutes), it was observed that the distinction between positive and negative reactions was much more apparent, providing for a quick naked-eye determination (Fig. 5b left). However, the CTV assays produced a less distinct color transition than the COX assays in the CTV-positive samples, likely due to the low initial copies of CTV in the sample. However, there was no increased distinction in fluorescent intensity between positive and negative reactions (Fig. 5b right). This study proved the quick sample preparation protocol integrated with lyophilized RT-LAMP assays was viable for in-greenhouse tests. This protocol has great potential to be adapted for the diagnosis of various plant diseases in the field.

**Fig. 5.**
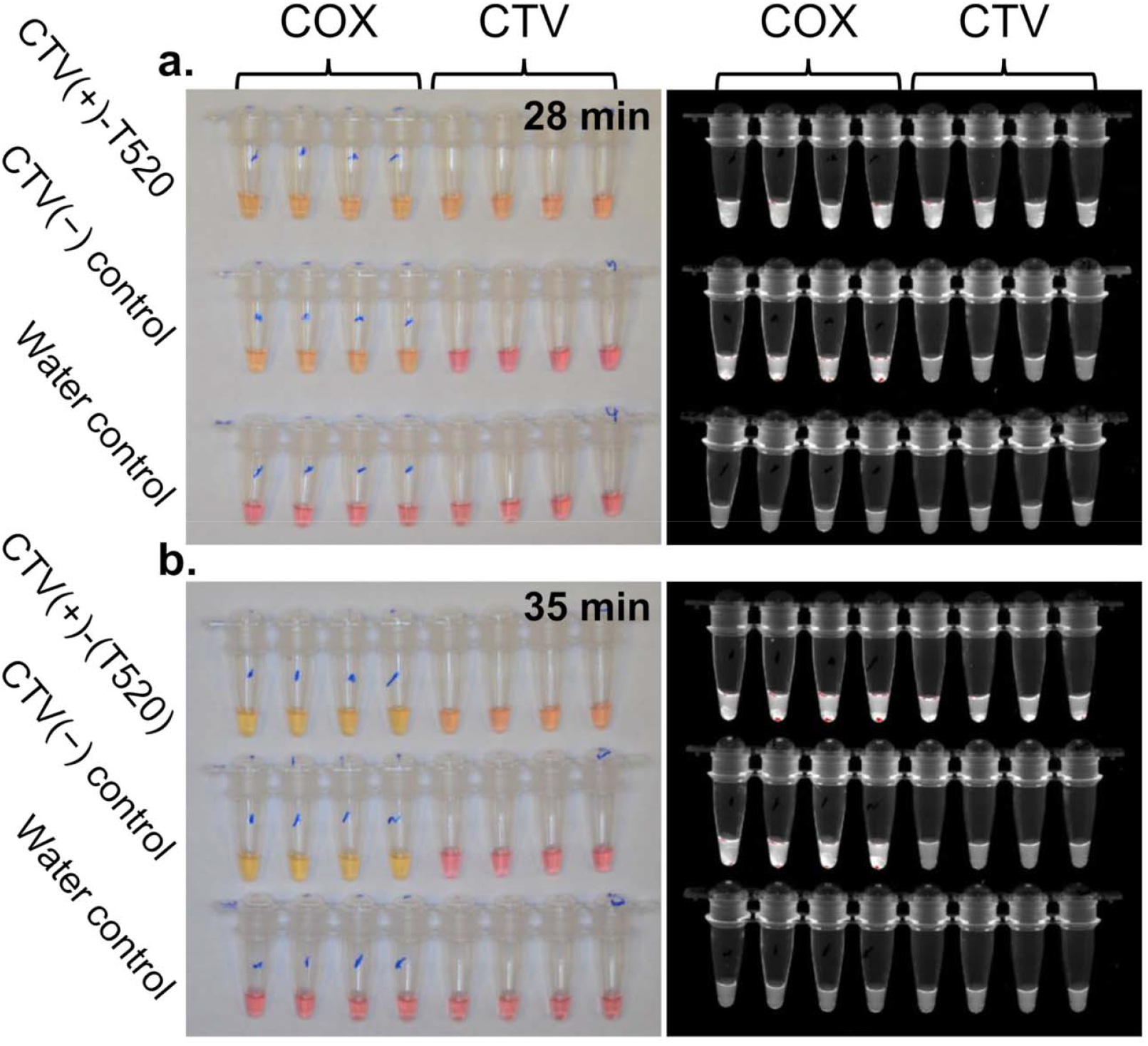
The evaluation of in-greenhouse RT-LAMP assays for CTV and COX detection. (a) Colorimetric assays (left) and fluorescence readout (right) after a 28-minute incubation at 65°C. (b) Colorimetric assays (left) and fluorescence readout (right) after a 35-minute incubation at 65°C.

## 4 Conclusion

In this study, a sensitive platform combining quick sample preparation and colorimetric RT-LAMP assays was demonstrated. Specifically, a previously developed protocol using micro-homogenizer and cellulose paper for quick preparation of nucleic acids was adopted by combining with RT-LAMP assays for visualized detection of CTV in a greenhouse. The RT-LAMP assays were optimized for the detection of CTV and COX in terms of concentrations of primers and reduction of false positives. Low detection limits were also achieved for CTV (43 copies/μL equivalent to 86 copies/mg of tissue) and COX (5 copies/μL equivalent to 10 copies/mg of tissue) RT-LAMP assays, suggesting an assay sensitivity comparable to qPCR-based techniques. Further, the RT-LAMP assays showed no false negatives or false positive

(Table S4). Ultimately, this platform combining lyophilized RT-LAMP reaction mix was successfully utilized in a greenhouse for onsite CTV detection, where the endpoint results were ready-to-read without a false response after 35 minutes of incubation at 65°C.

The proposed platform is a potential option for in-field plant disease diagnosis, where time-consuming procedures required in current practice, such as specimen collection, packaging, and submission steps, would no longer be needed. This technique expedites current disease diagnosis practice by saving significant time and cost for disease management, allowing the farmers and growers to carry out sample preparation and subsequent analysis on the spot. Overall, the successful demonstration of in-greenhouse RT-LAMP assays with a lyophilized reaction mix further expands the possibility of developing a sample-to-answer system to satisfy the increasing demands of in-field plant disease diagnosis.

## CRediT authorship contribution statement

**C.-W. L**.: Conceptualization, Methodology, Validation, Formal analysis, Investigation, Writing – original draft, Writing – Review & Editing, Visualization. **S. B**.: Conceptualization, Methodology, Resources, Writing – Review & Editing, Supervision, Funding acquisition. **B. K**.: Conceptualization, Methodology, Writing – original draft, Writing – Review & Editing, Visualization. **M. L. K**.: Methodology and Editing. **G. V**.: Conceptualization, Resources, Writing – Review & Editing, Supervision, Project administration, Funding acquisition. **H. T**.: Conceptualization, Writing – Review & Editing, Supervision, Project administration, Funding acquisition.

## Acknowledgments

The authors thank Dr. Arunabha Mitra for providing access to his citrus plants in Agricultural Operations (AgOPs) at the University of California, Riverside.

## Data Availability Statement

The data supporting the findings of this study are available from the corresponding author upon reasonable request.

## References

[1] S. He, K.M.C. Krainer, Pandemics of people and plants: which is the greater threat to food security?, Molecular plant 13(7) (2020) 933–934.

[2] B.K. Singh, M. Delgado-Baquerizo, E. Egidi, E. Guirado, J.E. Leach, H. Liu, P. Trivedi, Climate change impacts on plant pathogens, food security and paths forward, Nature Reviews Microbiology 21(10) (2023) 640–656.

[3] S. Tatineni, G.L. Hein, Plant viruses of agricultural importance: Current and future perspectives of virus disease management strategies, Phytopathology® 113(2) (2023) 117–141.

[4] World Food Programme, Global Report on Food Crises (GRFC) 2024, 2024. https://www.fsinplatform.org/report/global-report-food-crises-2024/. (Accessed August 4, 2024).

[5] J.B. Ristaino, P.K. Anderson, D.P. Bebber, K.A. Brauman, N.J. Cunniffe, N.V. Fedoroff, C. Finegold, K.A. Garrett, C.A. Gilligan, C.M. Jones, The persistent threat of emerging plant disease pandemics to global food security, Proceedings of the National Academy of Sciences 118(23) (2021) e2022239118.

[6] S. Atta, M. Cao, U.u.d. Umar, Y. Zhou, F. Yang, C. Zhou, Biological indexing and genetic analysis of Citrus tristeza virus in Pakistan, Journal of General Plant Pathology 83 (2017) 382–389.

[7] S. Hajeri, N. Killiny, C. El-Mohtar, W.O. Dawson, S. Gowda, Citrus tristeza virus-based RNAi in citrus plants induces gene silencing in Diaphorina citri, a phloem-sap sucking insect vector of citrus greening disease (Huanglongbing), Journal of biotechnology 176 (2014) 42–49.

[8] A. Karasev, V. Boyko, S. Gowda, O. Nikolaeva, M. Hilf, E. Koonin, C. Niblett, K. Cline, D. Gumpf, R. Lee, Complete sequence of the citrus tristeza virus RNA genome, Virology 208(2) (1995) 511–520.

[9] M. Ayllón, C. López, J. Navas-Castillo, S. Garnsey, J. Guerri, R. Flores, P. Moreno, Polymorphism of the 5′ terminal region of Citrus tristeza virus (CTV) RNA: incidence of three sequence types in isolates of different origin and pathogenicity, Archives of Virology 146 (2001) 27–40.

[10] K.K. Biswas, S. Palchoudhury, S.K. Sharma, B. Saha, S. Godara, D.K. Ghosh, M.L. Keremane, Analyses of 3’half genome of citrus tristeza virus reveal existence of distinct virus genotypes in citrus growing regions of India, VirusDisease 29 (2018) 308–315.

[11] W.B. Walker, M.L. Allen, Expression and RNA interference of salivary polygalacturonase genes in the tarnished plant bug, Lygus lineolaris, Journal of Insect Science 10(1) (2010) 173.

[12] J. Wang, O. Bozan, S.-J. Kwon, T. Dang, T. Rucker, R.K. Yokomi, R.F. Lee, S.Y. Folimonova, R.R. Krueger, J. Bash, Past and future of a century old Citrus tristeza virus collection: a California citrus germplasm tale, Frontiers in Microbiology 4 (2013) 366.

[13] J.M. Bové, Huanglongbing: a destructive, newly-emerging, century-old disease of citrus, Journal of plant pathology (2006) 7–37.

[14] California Department of Food & Agriculture, Sample Preparation and Submission Guideline. https://www.cdfa.ca.gov/plant/ppd/PDF/Submission_guidelines_Plant_Pathology.pdf. (Accessed August 03, 2024).

[15] T. Dang, F. Osman, J. Wang, T. Rucker, S. Bodaghi, S.-h. Tan, D. Pagliaccia, I. Lavagi-Craddock, G. Vidalakis, High-throughput RNA extraction from citrus tissues for the detection of viroids, Viroids: Methods and Protocols (2022) 57–64.

[16] S.A. Miller, F.D. Beed, C.L. Harmon, Plant disease diagnostic capabilities and networks, Annual review of phytopathology 47 (2009) 15–38.

[17] QIAGEN, RNeasy Plant Mini Kit for RNA Extraction. https://www.qiagen.com/us/products/discovery-and-translational-research/dna-rna-purification/rna-purification/total-rna/rneasy-plant-mini-kit. (Accessed August 28, 2024).

[18] QIAGEN, DNeasy Plant Kits. https://www.qiagen.com/us/products/discovery-and-translational-research/dna-rna-purification/dna-purification/genomic-dna/dneasy-plant-kits. (Accessed August 28, 2024).

[19] J. Singh, D. Cobb-Smith, E. Higgins, A. Khan, Comparative evaluation of lateral flow immunoassays, LAMP, and quantitative PCR for diagnosis of fire blight in apple orchards, Journal of Plant Pathology 103 (2021) 131–142.

[20] A. Kanapiya, U. Amanbayeva, Z. Tulegenova, A. Abash, S. Zhangazin, K. Dyussembayev, G. Mukiyanova, Recent advances and challenges in plant viral diagnostics, Frontiers in Plant Science 15 (2024) 1451790.

[21] M. Kumar, S.R. Kavalappara, T. McAvoy, S. Hutton, A.M. Simmons, S. Bag, Association of Tomato Chlorosis Virus Complicates the Management of Tomato Yellow Leaf Curl Virus in Cultivated Tomato (Solanum lycopersicum) in the Southern United States, Horticulturae 9(8) (2023) 948.

[22] S. Schneider, E. Jung, V. Queloz, J.B. Meyer, D. Rigling, Detection of pine needle diseases caused by Dothistroma septosporum, Dothistroma pini and Lecanosticta acicola using different methodologies, Forest Pathology 49(2) (2019) e12495.

[23] R. Paul, A.C. Saville, J.C. Hansel, Y. Ye, C. Ball, A. Williams, X. Chang, G. Chen, Z. Gu, J.B. Ristaino, Extraction of plant DNA by microneedle patch for rapid detection of plant diseases, ACS nano 13(6) (2019) 6540–6549.

[24] C.-W. Liu, B. Kalish, S. Bodaghi, G. Vidalakis, H. Tsutsui, A 3D-printed handheld device for quick citrus tissue lysis and nucleic acid extraction, bioRxiv (2024) 2024.08. 26.609775.

[25] N.J. Haveman, A.C. Schuerger, P.-L. Yu, M. Brown, R. Doebler, A.-L. Paul, R.J. Ferl,Advancing the automation of plant nucleic acid extraction for rapid diagnosis of plant diseases in space, Frontiers in Plant Science 14 (2023) 1194753.

[26] C.-W. Liu, S. Bodaghi, G. Vidalakis, H. Tsutsui, Quick Plant Sample Preparation Methods Using a Micro-Homogenizer for the Detection of Multiple Citrus Pathogens, Chemosensors 12(6) (2024) 105.

[27] C.-W. Liu, H. Tsutsui, Sample–to-answer sensing technologies for nucleic acid preparation and detection in the field, SLAS Technology 28(5) (2023) 302–323.

[28] Y. Zou, M.G. Mason, Y. Wang, E. Wee, C. Turni, P.J. Blackall, M. Trau, J.R. Botella, Nucleic acid purification from plants, animals and microbes in under 30 seconds, PLoS biology 15(11) (2017) e2003916.

[29] D.M. Mathews, S. Bodaghi, J.A. Heick, J.A. Dodds, Detection of Avocado Sunblotch and Other Viroids Using RNA Filter Paper Capture and RT-PCR, Viroids: Methods and Protocols (2022) 219–233.

[30] P. Li, M. Li, D. Yue, H. Chen, SolidlJphase extraction methods for nucleic acid separation. A review, Journal of Separation Science 45(1) (2022) 172–184.

[31] T.J. Moehling, G. Choi, L.C. Dugan, M. Salit, R.J. Meagher, LAMP diagnostics at the point-of-care: emerging trends and perspectives for the developer community, Expert Review of Molecular Diagnostics 21(1) (2021) 43–61.

[32] T. Notomi, Y. Mori, N. Tomita, H. Kanda, Loop-mediated isothermal amplification (LAMP): principle, features, and future prospects, Journal of microbiology 53(1) (2015) 1–5.

[33] T. Shymanovich, A.C. Saville, R. Paul, Q. Wei, J.B. Ristaino, Rapid detection of viral, bacterial, fungal, and oomycete pathogens on tomato with microneedles, LAMP on a microfluidic chip, and smartphone device, Phytopathology (ja) (2024).

[34] R. Paul, E. Ostermann, Y. Chen, A.C. Saville, Y. Yang, Z. Gu, A.E. Whitfield, J.B. Ristaino, Q. Wei, Integrated microneedle-smartphone nucleic acid amplification platform for in-field diagnosis of plant diseases, Biosensors and Bioelectronics 187 (2021) 113312.

[35] F. Osman, E. Hodzic, S.-J. Kwon, J. Wang, G. Vidalakis, Development and validation of a multiplex reverse transcription quantitative PCR (RT-qPCR) assay for the rapid detection of Citrus tristeza virus, Citrus psorosis virus, and Citrus leaf blotch virus, Journal of Virological Methods 220 (2015) 64–75.

[36] New England Biolabs, WarmStart Colorimetric LAMP 2X Master Mix Typical LAMP Protocol (M1800). https://www.neb.com/en-us/protocols/2016/08/15/warmstart-colorimetric-lamp-2x-master-mix-typical-lamp-protocol-m1800. (Accessed August 2, 2024).

[37] X. Song, F.J. Coulter, M. Yang, J.L. Smith, F.G. Tafesse, W.B. Messer, J.H. Reif, A lyophilized colorimetric RT-LAMP test kit for rapid, low-cost, at-home molecular testing of SARS-CoV-2 and other pathogens, Scientific Reports 12(1) (2022) 7043.

[38] V. Selvaraj, Y. Maheshwari, S. Hajeri, R. Yokomi, A rapid detection tool for VT isolates of Citrus tristeza virus by immunocapture-reverse transcriptase loop-mediated isothermal amplification assay, PLoS One 14(9) (2019) e0222170.

[39] Y. Shao, S. Zhu, C. Jin, F. Chen, Development of multiplex loop-mediated isothermal amplification-RFLP (mLAMP-RFLP) to detect Salmonella spp. and Shigella spp. in milk, International journal of food microbiology 148(2) (2011) 75–79.

[40] D.-G. Wang, J.D. Brewster, M. Paul, P.M. Tomasula, Two methods for increased specificity and sensitivity in loop-mediated isothermal amplification, Molecules 20(4) (2015) 6048–6059.

[41] N.K. Jain, I. Roy, Trehalose and Protein Stability, Current Protocols in Protein Science 59(1) (2010) 4.9.1-4.9.12.

[42] Y. Zhang, G. Ren, J. Buss, A.J. Barry, G.C. Patton, N.A. Tanner, Enhancing Colorimetric Loop-mediated Isothermal Amplification Speed and Sensitivity with Guanidine Chloride, BioTechniques 69(3) (2020) 178–185.

