## Supplementary Information for "Rapid colorimetric detection of *citrus tristeza virus* combining portable sample preparation and reverse transcription-loop mediated isothermal amplification"


Table S1. Alignment of coat protein gene sequences of Citrus tristeza virus using BLAST program of the NCBI (https://blast.ncbi.nlm.nih.gov/). Sequences used for LAMP primers are shown.

DownloadNextPreviousFirst Range

Query range 8: 421 to 480 → LAMP-F3 (CTV) → LAMP-F2 (CTV) LAMP-LFc (CTV)

| | | | |
|---|---|---|---|
| Query | 421 | AGAACTAACGATGCTCTTTACTTAGCCTTTTGTAGACAGAACCGCAATTTGAGTTATGGC | 480 |
| AJ518842.1 | 421 | ............................................................ | 480 |
| KU365761.1 | 421 | ............................................................ | 480 |
| KU365760.1 | 421 | ............................................................ | 480 |
| MT833084.1 | 421 | ............................................................ | 480 |
| MT833082.1 | 421 | ............................................................ | 480 |
| MT833081.1 | 421 | ............................................................ | 480 |
| MT833071.1 | 421 | ............................................................ | 480 |
| MT833060.1 | 421 | ............................................................ | 480 |
| MT833059.1 | 421 | ............................................................ | 480 |
| MT833012.1 | 421 | ............................................................ | 480 |
| MT833083.1 | 421 | ............................................................ | 480 |
| MT833074.1 | 421 | ............................................................ | 480 |
| AM746968.2 | 421 | ............................................................ | 480 |
| OK082186.1 | 421 | ............................................................ | 480 |
| MT833212.1 | 421 | ............................................................ | 480 |
| MT833211.1 | 421 | ............................................................ | 480 |
| MT833210.1 | 421 | ............................................................ | 480 |
| MT833209.1 | 421 | ............................................................ | 480 |
| MT833207.1 | 421 | ............................................................ | 480 |
| OR122727.1 | 16441 | ............................................................ | 16500 |
| MT833213.1 | 421 | ............................................................ | 480 |
| MT833208.1 | 421 | ............................................................ | 480 |
| MT833206.1 | 421 | ............................................................ | 480 |
| EU660931.1 | 421 | ......................................A..................... | 480 |
| MT833205.1 | 421 | ...........................................A................ | 480 |
| MT833204.1 | 421 | ...........................................A................ | 480 |
| EU660932.1 | 421 | ............................................................ | 480 |
| KU365759.1 | 421 | ..................C......................................... | 480 |
| OK082164.1 | 421 | ............................................................ | 480 |
| OK082169.1 | 421 | ............................................................ | 480 |
| OK082167.1 | 421 | ............................................................ | 480 |
| KF962599.1 | 421 | ............................................................ | 480 |
| MW201832.1 | 421 | ............................................................ | 480 |
| EU660910.1 | 421 | ............................................................ | 480 |
| EU660909.1 | 421 | ............................................................ | 480 |
| MZ330116.1 | 16536 | ............................................................ | 16595 |
| MT833135.1 | 421 | ............................................................ | 480 |
| MT833133.1 | 421 | ............................................................ | 480 |

```
MT833130.1  421    ............................................................  480
MT833128.1  421    ............................................................  480
GU983388.1  421    ............................................................  480
AY995567.1  4705   ............................................................  4764
AY995566.1  4702   ............................................................  4761
AY995565.1  4702   ............................................................  4761
AY995564.1  4702   ............................................................  4761
AY995563.1  4705   ............................................................  4764
AY995562.1  4705   ............................................................  4764
AM406802.1  421    ............................................................  480
MZ648331.1  16534  ............................................................  16593
KF962600.1  421    .......................G....................................  480
KF962598.1  421    ............................................................  480
MT833129.1  421    ............................................................  480
KC748391.1  16536  ............................................................  16595
KC841807.1  421    ............................................................  480
KC841796.1  421    ............................................................  480
KC841795.1  421    ............................................................  480
KC841792.1  421    ............................................................  480
KC841783.1  421    ............................................................  480
HF947334.1  421    ............................................................  480
FR856889.1  421    ............................................................  480
GU983386.1  421    ............................................................  480
GQ424345.1  421    ............................................................  480
FN661497.1  421    ............................................................  480
EU878378.1  595    ............................................................  654
AM406803.1  421    ............................................................  480
Y18420.1    16536  ............................................................  16595
MK049162.1  421    ............................................................  480
GQ424348.1  421    ............................................................  480
EU878384.1  570    ............................................................  629
MK779711.1  16537  ............................................................  16596
MH279618.1  16536  ............................................................  16595
KP284576.1  421    ............................................................  480
MT833189.1  421    ............................................................  480
MT833187.1  421    ............................................................  480
MT833170.1  421    ............................................................  480
MT833138.1  421    ............................................................  480
MT833134.1  421    ............................................................  480
MT833132.1  421    ............................................................  480
MT833131.1  421    ............................................................  480
KC841811.1  421    ............................................................  480
KC841805.1  421    ............................................................  480
KC841786.1  421    ...........C................................................  480
HF947337.1  421    ............................................................  480
KC517491.1  16530  ............................................................  16589
KC517490.1  16527  ............................................................  16586
JQ339726.1  421    ............................................................  480
FR871881.1  421    ............................................................  480
GU983384.1  421    ............................................................  480
FN661494.1  421    ............................................................  480
GQ424358.1  421    ............................................................  480
GQ424353.1  421    ............................................................  480
EU579374.1  421    ............................................................  480
EU937520.1  16537  ............................................................  16596
AM746969.1  421    ............................................................  480
AF220503.1  421    ............................................................  480
AF260651.1  16536  ............................................................  16595
OP006461.1  421    ............................................................  480
MZ670756.1  421    ............................................................  480
MT833188.1  421    ............................................................  480
KF196264.1  421    ............................................................  480
```

DownloadNextPreviousFirst Range

```
Query range 9: 481 to 540    ←        LAMP-F1 (CTV)  ←→ LAMP-B1c (CTV) (ATCT >> ATTT)
Query       481    GGACGTCCGCTAGATGCAGGGATTCCGGCTGGGTATCATTATCTGTGTGCAGATTTCTTG  540
AJ518842.1  481    ............................................................  540
                                                           →  PCR-F1, F2 (CTV)
KU365761.1  481    ............................................................  540
KU365760.1  481    ............................................................  540
```

| | | | |
|---|---|---|---|
| MT833084.1 | 481 | ............................................................ | 540 |
| MT833082.1 | 481 | ............................................................ | 540 |
| MT833081.1 | 481 | ............................................................ | 540 |
| MT833071.1 | 481 | ............................................................ | 540 |
| MT833060.1 | 481 | ............................................................ | 540 |
| MT833059.1 | 481 | ............................................................ | 540 |
| MT833012.1 | 481 | ............................................................ | 540 |
| MT833083.1 | 481 | ............................................................ | 540 |
| MT833074.1 | 481 | ...............................T............................ | 540 |
| AM746968.2 | 481 | ..........................................T................. | 540 |
| OK082186.1 | 481 | ............................................................ | 540 |
| MT833212.1 | 481 | ............................................................ | 540 |
| MT833211.1 | 481 | ............................................................ | 540 |
| MT833210.1 | 481 | ............................................................ | 540 |
| MT833209.1 | 481 | ............................................................ | 540 |
| MT833207.1 | 481 | ............................................................ | 540 |
| OR122727.1 | 16501 | ............................................................ | 16560 |
| MT833213.1 | 481 | ............................................................ | 540 |
| MT833208.1 | 481 | ..............C............................................. | 540 |
| MT833206.1 | 481 | ............................................................ | 540 |
| EU660931.1 | 481 | ............................................................ | 540 |
| MT833205.1 | 481 | ............................................................ | 540 |
| MT833204.1 | 481 | ............................................................ | 540 |
| EU660932.1 | 481 | .............................C.............................. | 540 |
| KU365759.1 | 481 | ............................................................ | 540 |
| OK082164.1 | 481 | ............................................................ | 540 |
| OK082169.1 | 481 | .............................C.............................. | 540 |
| OK082167.1 | 481 | ............................................................ | 540 |
| KF962599.1 | 481 | ..........................................T................. | 540 |
| MW201832.1 | 481 | .............................C.............................. | 540 |
| EU660910.1 | 481 | ............................................................ | 540 |
| EU660909.1 | 481 | .......................................C.................... | 540 |
| MZ330116.1 | 16596 | ..........................................T................. | 16655 |
| MT833135.1 | 481 | ............................................................ | 540 |
| MT833133.1 | 481 | ............................................................ | 540 |
| MT833130.1 | 481 | ............................................................ | 540 |
| MT833128.1 | 481 | ............................................................ | 540 |
| GU983388.1 | 481 | ..........................................T................. | 540 |
| AY995567.1 | 4765 | ..........................................T................. | 4824 |
| AY995566.1 | 4762 | ..........................................T................. | 4821 |
| AY995565.1 | 4762 | ..........................................T................. | 4821 |
| AY995564.1 | 4762 | ..........................................T................. | 4821 |
| AY995563.1 | 4765 | ..........................................T................. | 4824 |
| AY995562.1 | 4765 | ..........................................T................. | 4824 |
| AM406802.1 | 481 | ..........................................T................. | 540 |
| MZ648331.1 | 16594 | .........................................CT................. | 16653 |
| KF962600.1 | 481 | ..........................................T................. | 540 |
| KF962598.1 | 481 | ..........................................T................. | 540 |
| MT833129.1 | 481 | ............................................................ | 540 |
| KC748391.1 | 16596 | ..........................................T................. | 16655 |
| KC841807.1 | 481 | ..........................................T................. | 540 |
| KC841796.1 | 481 | ..........................................T................. | 540 |
| KC841795.1 | 481 | ..........................................T................. | 540 |
| KC841792.1 | 481 | ..........................................T................. | 540 |
| KC841783.1 | 481 | ..........................................T................. | 540 |
| HF947334.1 | 481 | ..........................................T................. | 540 |
| FR856889.1 | 481 | ..........................................T................. | 540 |
| GU983386.1 | 481 | ..................C.......................T................. | 540 |
| GQ424345.1 | 481 | ..........................................T................. | 540 |
| FN661497.1 | 481 | ..........................................T................. | 540 |
| EU878378.1 | 655 | ..........................................T................. | 714 |
| AM406803.1 | 481 | ..........................................T................. | 540 |
| Y18420.1 | 16596 | ..........................................T................. | 16655 |
| MK049162.1 | 481 | ..........................................T................. | 540 |
| GQ424348.1 | 481 | ..........................................T................. | 540 |
| EU878384.1 | 630 | ..........................................T................. | 689 |
| MK779711.1 | 16597 | ..........................................T................. | 16656 |
| MH279618.1 | 16596 | ..........................................T................. | 16655 |
| KP284576.1 | 481 | ..........................................T..............C.. | 540 |
| MT833189.1 | 481 | ........T.................................T................. | 540 |
| MT833187.1 | 481 | ........T.................................T................. | 540 |

```
MT833170.1  481    ..........................................T.................  540
MT833138.1  481    ............................................................  540
MT833134.1  481    ............................................................  540
MT833132.1  481    ............................................................  540
MT833131.1  481    ............................................................  540
KC841811.1  481    ..........................................T.................  540
KC841805.1  481    ..........................................T.................  540
KC841786.1  481    ..........................................T.................  540
HF947337.1  481    ..........................................T.................  540
KC517491.1  16590  ..........................................T.................  16649
KC517490.1  16587  ..........................................T.................  16646
JQ339726.1  481    ..........................................T.................  540
FR871881.1  481    ..........................................T.................  540
GU983384.1  481    ..........................................T.................  540
FN661494.1  481    ..........................................T.................  540
GQ424358.1  481    ..........................................T.................  540
GQ424353.1  481    ..........................................T.................  540
EU579374.1  481    ........A..........................C........................  540
EU937520.1  16597  ..........................................T.................  16656
AM746969.1  481    ..........................................T.................  540
AF220503.1  481    ..........................................T.................  540
AF260651.1  16596  ..........................................T.................  16655
OP006461.1  481    ..........................................T.................  540
MZ670756.1  481    ..........................................T.................  540
MT833188.1  481    ........T.................................T.................  540
KF196264.1  481    ..........................................T.................  540
```

DownloadNextPreviousFirst Range

Query range 10: 541 to 600 → LAMP-LB (CTV) LAMP-B2c (CTV) ←

```
Query       541    ACCGGAGCTGGCTTGACTGATTTAGAATGTGCTGTGTACATACAAGCTAAAGAACAATTG  600
AJ518842.1  541    ............................................................  600
KU365761.1  541    ............................................................  600
KU365760.1  541    ..................................G.........................  600
MT833084.1  541    ............................................................  600
MT833082.1  541    ............................................................  600
MT833081.1  541    ............................................................  600
MT833071.1  541    ............................................................  600
MT833060.1  541    ............................................................  600
MT833059.1  541    ............................................................  600
MT833012.1  541    ............................................................  600
MT833083.1  541    ............................................................  600
MT833074.1  541    ..........................................................G.  600
AM746968.2  541    ............................................................  600
OK082186.1  541    ............................................................  600
MT833212.1  541    ............................................................  600
MT833211.1  541    ........................................................T...  600
MT833210.1  541    ............................................................  600
MT833209.1  541    ............................................................  600
MT833207.1  541    ............................................................  600
OR122727.1  16561  ............................................................  16620
MT833213.1  541    ............................................................  600
MT833208.1  541    ............................................................  600
MT833206.1  541    ............................................................  600
EU660931.1  541    ............................................................  600
MT833205.1  541    ..........................G.................................  600
MT833204.1  541    ..........................G.................................  600
EU660932.1  541    ..................................C.........................  600
KU365759.1  541    .............................G....G.........................  600
OK082164.1  541    ............................................................  600
OK082169.1  541    ............................................................  600
OK082167.1  541    ............................................................  600
KF962599.1  541    ............................................................  600
MW201832.1  541    ............................................................  600
EU660910.1  541    G...........................................................  600
EU660909.1  541    ............................................................  600
MZ330116.1  16656  ............................................................  16715
MT833135.1  541    ............................................................  600
MT833133.1  541    ............................................................  600
MT833130.1  541    ............................................................  600
MT833128.1  541    ............................................................  600
```

```
GU983388.1  541    ............................................................  600
AY995567.1  4825   ............................................................  4884
AY995566.1  4822   ............................................................  4881
AY995565.1  4822   ............................................................  4881
AY995564.1  4822   ............................................................  4881
AY995563.1  4825   ............................................................  4884
AY995562.1  4825   ............................................................  4884
AM406802.1  541    ............................................................  600
MZ648331.1  16654  ............................................................  16713
KF962600.1  541    ............................................................  600
KF962598.1  541    ............................................................  600
MT833129.1  541    ............................................................  600
KC748391.1  16656  ............................................................  16715
KC841807.1  541    .................C..........................................  600
KC841796.1  541    ............................................................  600
KC841795.1  541    ............................................................  600
KC841792.1  541    ............................................................  600
KC841783.1  541    ............................................................  600
HF947334.1  541    ............................................................  600
FR856889.1  541    ............................................................  600
GU983386.1  541    ............................................................  600
GQ424345.1  541    ............................................................  600
FN661497.1  541    ............................................................  600
EU878378.1  715    ............................................................  774
AM406803.1  541    ............................................................  600
Y18420.1    16656  ............................................................  16715
MK049162.1  541    ............................................................  600
GQ424348.1  541    ............................................................  600
EU878384.1  690    ............................................................  749
MK779711.1  16657  ............................................................  16716
MH279618.1  16656  ............................................................  16715
KP284576.1  541    ............................................................  600
MT833189.1  541    ............................................................  600
MT833187.1  541    ............................................................  600
MT833170.1  541    ............................................................  600
MT833138.1  541    ....A.......................................................  600
MT833134.1  541    ....A.......................................................  600
MT833132.1  541    ............................................................  600
MT833131.1  541    ....A.......................................................  600
KC841811.1  541    ............................................................  600
KC841805.1  541    ............................................................  600
KC841786.1  541    ............................................................  600
HF947337.1  541    ........................................................G...  600
KC517491.1  16650  ............................................................  16709
KC517490.1  16647  ............................................................  16706
JQ339726.1  541    ............................................................  600
FR871881.1  541    ..........C.................................................  600
GU983384.1  541    ............................................................  600
FN661494.1  541    ............................................................  600
GQ424358.1  541    ............................................................  600
GQ424353.1  541    ............................................................  600
EU579374.1  541    ............................................................  600
EU937520.1  16657  ............................................................  16716
AM746969.1  541    ............................................................  600
AF220503.1  541    ............................................................  600
AF260651.1  16656  ............................................................  16715
OP006461.1  541    ............................................................  600
MZ670756.1  541    ............................................................  600
MT833188.1  541    ............................................................  600
KF196264.1  541    ............................................................  600
```

DownloadNextPreviousFirst Range

Query range 11: 601 to 660 LAMP-B3c (CTV) ← PCR-Rc (CTV) ←

```
Query       601    TTGAAAAAGCGAGGGGCTGATGAGGTTGTAGTTACTAATGTCAGGCAGCTTGGGAAATTT  660
                         → PCR-Probe (CTV)
AJ518842.1  601    ............................................................  660
KU365761.1  601    ............................................................  660
KU365760.1  601    ............................................................  660
MT833084.1  601    ...........................................................C  660
MT833082.1  601    ...........................................................C  660
```

| | | | |
|---|---|---|---|
| MT833081.1 | 601 | ...........................................................C | 660 |
| MT833071.1 | 601 | ...........................................................C | 660 |
| MT833060.1 | 601 | ...........................................................C | 660 |
| MT833059.1 | 601 | ...........................................................C | 660 |
| MT833012.1 | 601 | ...........................................................C | 660 |
| MT833083.1 | 601 | ...........................................................C | 660 |
| MT833074.1 | 601 | ...................G................................C......C | 660 |
| AM746968.2 | 601 | .............................G.............................. | 660 |
| OK082186.1 | 601 | ...........................................................C | 660 |
| MT833212.1 | 601 | ...........................................................C | 660 |
| MT833211.1 | 601 | ...........................................................C | 660 |
| MT833210.1 | 601 | ...........................................................C | 660 |
| MT833209.1 | 601 | ...........................................................C | 660 |
| MT833207.1 | 601 | ...........................................................C | 660 |
| OR122727.1 | 16621 | ............................................................ | 16680 |
| MT833213.1 | 601 | ....................................................A......C | 660 |
| MT833208.1 | 601 | ...........................................................C | 660 |
| MT833206.1 | 601 | ...........................................................C | 660 |
| EU660931.1 | 601 | ...........................................................C | 660 |
| MT833205.1 | 601 | ...........................................................C | 660 |
| MT833204.1 | 601 | ...........................................................C | 660 |
| EU660932.1 | 601 | ...........................................................C | 660 |
| KU365759.1 | 601 | ........................................G................... | 660 |
| OK082164.1 | 601 | ...........................................................C | 660 |
| OK082169.1 | 601 | .....G.....................................................C | 660 |
| OK082167.1 | 601 | ...........................................................C | 660 |
| KF962599.1 | 601 | ............................................................ | 660 |
| MW201832.1 | 601 | .....G.....................................................C | 660 |
| EU660910.1 | 601 | ...........................................................C | 660 |
| EU660909.1 | 601 | ...........................................................C | 660 |
| MZ330116.1 | 16716 | ............................................................ | 16775 |
| MT833135.1 | 601 | ...........................................................C | 660 |
| MT833133.1 | 601 | ...........................................................C | 660 |
| MT833130.1 | 601 | ...........................................................C | 660 |
| MT833128.1 | 601 | ...........................................................C | 660 |
| GU983388.1 | 601 | ............................................................ | 660 |
| AY995567.1 | 4885 | ............................................................ | 4944 |
| AY995566.1 | 4882 | ............................................................ | 4941 |
| AY995565.1 | 4882 | ............................................................ | 4941 |
| AY995564.1 | 4882 | ............................................................ | 4941 |
| AY995563.1 | 4885 | ............................................................ | 4944 |
| AY995562.1 | 4885 | ............................................................ | 4944 |
| AM406802.1 | 601 | ............................................................ | 660 |
| MZ648331.1 | 16714 | ............................................................ | 16773 |
| KF962600.1 | 601 | ............................................................ | 660 |
| KF962598.1 | 601 | ............................................................ | 660 |
| MT833129.1 | 601 | ....................................................A......C | 660 |
| KC748391.1 | 16716 | ............................................................ | 16775 |
| KC841807.1 | 601 | ............................................................ | 660 |
| KC841796.1 | 601 | ............................................................ | 660 |
| KC841795.1 | 601 | ....................C....................................... | 660 |
| KC841792.1 | 601 | ............................................................ | 660 |
| KC841783.1 | 601 | ............................................................ | 660 |
| HF947334.1 | 601 | ...........................................................C | 660 |
| FR856889.1 | 601 | ...........................................................C | 660 |
| GU983386.1 | 601 | ............................................................ | 660 |
| GQ424345.1 | 601 | ...........................................................C | 660 |
| FN661497.1 | 601 | ...........................................................C | 660 |
| EU878378.1 | 775 | ...........................................................C | 834 |
| AM406803.1 | 601 | ............................................................ | 660 |
| Y18420.1 | 16716 | ............................................................ | 16775 |
| MK049162.1 | 601 | ...........................................................C | 660 |
| GQ424348.1 | 601 | ...........................................N...............C | 660 |
| EU878384.1 | 750 | ...........................................................C | 809 |
| MK779711.1 | 16717 | ............................................................ | 16776 |
| MH279618.1 | 16716 | ............................................................ | 16775 |
| KP284576.1 | 601 | ............................................................ | 660 |
| MT833189.1 | 601 | ...........................................................C | 660 |
| MT833187.1 | 601 | ...........................................................C | 660 |
| MT833170.1 | 601 | ............................................................ | 660 |
| MT833138.1 | 601 | ...........................................................C | 660 |

```
MT833134.1   601     ...........................................................C  660
MT833132.1   601     .......................T............................A......C  660
MT833131.1   601     ...........................................................C  660
KC841811.1   601     ............................................................  660
KC841805.1   601     ............................................................  660
KC841786.1   601     ............................................................  660
HF947337.1   601     ...........................................................C  660
KC517491.1   16710   ............................................................  16769
KC517490.1   16707   ............................................................  16766
JQ339726.1   601     ...........................................................C  660
FR871881.1   601     ...........................................................C  660
GU983384.1   601     ............................................................  660
FN661494.1   601     ...........................................................C  660
GQ424358.1   601     ...........................................................C  660
GQ424353.1   601     ...........................................................C  660
EU579374.1   601     .................C........................T................C  660
EU937520.1   16717   ............................................................  16776
AM746969.1   601     ............................................................  660
AF220503.1   601     ........A...................................................  660
AF260651.1   16716   ............................................................  16775
OP006461.1   601     ...........................................................C  660
MZ670756.1   601     ............................................................  660
MT833188.1   601     ...........................................................C  660
KF196264.1   601     ..A........................................................C  660
```

DownloadNextPreviousFirst Range

Query range 12: 661 to 672

```
Query        661     AACACACGTTGA  672
AJ518842.1   661     ............  672
KU365761.1   661     ............  672
KU365760.1   661     ............  672
MT833084.1   661     ............  672
MT833082.1   661     ............  672
MT833081.1   661     ............  672
MT833071.1   661     ............  672
MT833060.1   661     ............  672
MT833059.1   661     ............  672
MT833012.1   661     ............  672
MT833083.1   661     ............  672
MT833074.1   661     ............  672
AM746968.2   661     ............  672
OK082186.1   661     ............  672
MT833212.1   661     ............  672
MT833211.1   661     ............  672
MT833210.1   661     ............  672
MT833209.1   661     ............  672
MT833207.1   661     ............  672
OR122727.1   16681   ............  16692
MT833213.1   661     ............  672
MT833208.1   661     ............  672
MT833206.1   661     ............  672
EU660931.1   661     ............  672
MT833205.1   661     ............  672
MT833204.1   661     ............  672
EU660932.1   661     ............  672
KU365759.1   661     ............  672
OK082164.1   661     ............  672
OK082169.1   661     ............  672
OK082167.1   661     ............  672
KF962599.1   661     ............  672
MW201832.1   661     ............  672
EU660910.1   661     ............  672
EU660909.1   661     ............  672
MZ330116.1   16776   ............  16787
MT833135.1   661     ............  672
MT833133.1   661     ............  672
MT833130.1   661     ............  672
MT833128.1   661     ............  672
GU983388.1   661     ............  672
AY995567.1   4945    ............  4956
```

| | | | |
|---|---|---|---|
| AY995566.1 | 4942 | ............ | 4953 |
| AY995565.1 | 4942 | ............ | 4953 |
| AY995564.1 | 4942 | ............ | 4953 |
| AY995563.1 | 4945 | ............ | 4956 |
| AY995562.1 | 4945 | ............ | 4956 |
| AM406802.1 | 661 | ............ | 672 |
| MZ648331.1 | 16774 | ............ | 16785 |
| KF962600.1 | 661 | ............ | 672 |
| KF962598.1 | 661 | ............ | 672 |
| MT833129.1 | 661 | ............ | 672 |
| KC748391.1 | 16776 | ............ | 16787 |
| KC841807.1 | 661 | ............ | 672 |
| KC841796.1 | 661 | ............ | 672 |
| KC841795.1 | 661 | ............ | 672 |
| KC841792.1 | 661 | ............ | 672 |
| KC841783.1 | 661 | ............ | 672 |
| HF947334.1 | 661 | ............ | 672 |
| FR856889.1 | 661 | ............ | 672 |
| GU983386.1 | 661 | ............ | 672 |
| GQ424345.1 | 661 | ............ | 672 |
| FN661497.1 | 661 | ............ | 672 |
| EU878378.1 | 835 | ............ | 846 |
| AM406803.1 | 661 | ............ | 672 |
| Y18420.1 | 16776 | ............ | 16787 |
| MK049162.1 | 661 | ........... | 671 |
| GQ424348.1 | 661 | ............ | 672 |
| EU878384.1 | 810 | .......... | 819 |
| MK779711.1 | 16777 | ............ | 16788 |
| MH279618.1 | 16776 | ............ | 16787 |
| KP284576.1 | 661 | ............ | 672 |
| MT833189.1 | 661 | ............ | 672 |
| MT833187.1 | 661 | ............ | 672 |
| MT833170.1 | 661 | ............ | 672 |
| MT833138.1 | 661 | ............ | 672 |
| MT833134.1 | 661 | ............ | 672 |
| MT833132.1 | 661 | ............ | 672 |
| MT833131.1 | 661 | ............ | 672 |
| KC841811.1 | 661 | ............ | 672 |
| KC841805.1 | 661 | ............ | 672 |
| KC841786.1 | 661 | ............ | 672 |
| HF947337.1 | 661 | ............ | 672 |
| KC517491.1 | 16770 | ............ | 16781 |
| KC517490.1 | 16767 | ............ | 16778 |
| JQ339726.1 | 661 | ............ | 672 |
| FR871881.1 | 661 | ............ | 672 |
| GU983384.1 | 661 | ............ | 672 |
| FN661494.1 | 661 | ............ | 672 |
| GQ424358.1 | 661 | ............ | 672 |
| GQ424353.1 | 661 | ............ | 672 |
| EU579374.1 | 661 | ............ | 672 |
| EU937520.1 | 16777 | ............ | 16788 |
| AM746969.1 | 661 | ............ | 672 |
| AF220503.1 | 661 | ............ | 672 |
| AF260651.1 | 16776 | ............ | 16787 |
| OP006461.1 | 661 | ............ | 672 |
| MZ670756.1 | 661 | ............ | 672 |
| MT833188.1 | 661 | ............ | 672 |
| KF196264.1 | 661 | ............ | 672 |

Table S2. Alignment of cytochrome oxidase gene of citrus using BLAST program of the NCBI (https://blast.ncbi.nlm.nih.gov/). Sequences used for LAMP primers are shown.

```
Query range 1: 1 to 60                                 → PCR-F (COX)              →
Query        5       CATTTTTGGATCACTTTTTTCGGGGTTAATCTGACCTTCTTTCCCATGCATTTCTTAGGG   64
NC_037463.1  478187  ............................................................   478128
PP894751.1   223460  ............................................................   223401
PP770439.1   349495  ............................................................   349436
```

```
PP035765.1   118137  ............................................................  118196
NC_086688.1  93446   ............................................................  93387
MN946545.1   249908  ............................................................  249967
NC_057143.1  436059  ............................................................  436118
NC_057142.1  436050  ............................................................  436109
KF933043.1   1                                   ................................  32
```

DownloadNextPreviousFirst Range

Query range 2: 61 to 120

```
                     LAMP-F3 (COX)        →  LAMP-F2 (COX)        LAMP-LFc (COX)  ←
Query        65      CTTTCGGGTATGCCACGTCGCATTCCAGATTATCCAGATGCTTACGCTGGATGGAATGCC  124
NC_037463.1  478127  ............................................................  478068
                                                        → PCR-Probe (COX)
PP894751.1   223400  ............................................................  223341
PP770439.1   349435  ............................................................  349376
PP035765.1   118197  ............................................................  118256
NC_086688.1  93386   ............................................................  93327
MN946545.1   249968  ............................................................  250027
NC_057143.1  436119  ............................................................  436178
NC_057142.1  436110  ............................................................  436169
KF933043.1   33      ............................................................  92
```

DownloadNextPreviousFirst Range

Query range 3: 121 to 180

```
                            LAMP-F1 (COX)       ←       →  LAMP-B1c (COX)
Query        125     CTTAGCAGTTTTGGCTCTTATATATCCGTAGTTGGGATTTGTTGTTTCTTCGTGGTCGTA  184
NC_037463.1  478067  .......................................C..C.................  478008
PP894751.1   223340  .......................................C..C.................  223281
PP770439.1   349375  .......................................C..C.................  349316
PP035765.1   118257  .......................................C..C.................  118316
NC_086688.1  93326   .......................................C..C.................  93267
MN946545.1   250028  .......................................C..C.................  250087
NC_057143.1  436179  .......................................C..C.................  436238
NC_057142.1  436170  .......................................C..C.................  436229
KF933043.1   93      .......................................C..C.................  152
```

DownloadNextPreviousFirst Range

Query range 4: 181 to 240

```
                     PCR-Rc  ←   →  LAMP-LB (COX)             LAMP-B2c (COX)  ←
Query        185     ACAATCACTTTAAGCAGTGGAAATAACAAAAGATGTGCTCCAAGTCCTTGGGCTGTTGAA  244
NC_037463.1  478007  ..........C.................................................  477948
PP894751.1   223280  ..........C.................................................  223221
PP770439.1   349315  ..........C.................................................  349256
PP035765.1   118317  ..........C.................................................  118376
NC_086688.1  93266   ..........C.................................................  93207
MN946545.1   250088  ..........C.................................................  250147
NC_057143.1  436239  ..........C.................................................  436298
NC_057142.1  436230  ..........C.................................................  436289
KF933043.1   153     ..........C.................................................  212
```

DownloadNextPreviousFirst Range

Query range 5: 241 to 300

```
                      LAMP-B3c (COX)  ←
Query        245     CAGAATTCAACCACACTGGAATGGATGGTACAAAGTCCTCCAGCTTTTCATACTTTTGGA  304
NC_037463.1  477947  ......C.........C...........................................  477888
PP894751.1   223220  ......C.........C...........................................  223161
PP770439.1   349255  ......C.........C...........................................  349196
PP035765.1   118377  ......C.........C...........................................  118436
NC_086688.1  93206   ......C.........C...........................................  93147
MN946545.1   250148  ......C.........C...........................................  250207
NC_057143.1  436299  ......C.........C...........................................  436358
NC_057142.1  436290  ......C.........C...........................................  436349
KF933043.1   213     ......C.........C...........................................  272
```

DownloadNextPreviousFirst Range

Query range 6: 301 to 360

```
Query        305     GAACTTCCAGCTATCAAGGAGACGAAAAGCTATGTGAAGTAAAAGAAGAAAAGATCGCCG  364
NC_037463.1  477887  ............................................................  477828
PP894751.1   223160  ............................................................  223101
PP770439.1   349195  ............................................................  349136
```

```
PP035765.1   118437  ............................................................  118496
NC_086688.1  93146   ............................................................  93087
MN946545.1   250208  ............................................................  250267
NC_057143.1  436359  ............................................................  436418
NC_057142.1  436350  ............................................................  436409
KF933043.1   273     ............................................................  332
```

Download Next Previous First Range

Query range 7: 361 to 394

```
Query        365     ACTGCTACTAAGAACCTAACAGAACATTnnnnnn  398
NC_037463.1  477827  ............................TC....  477794
PP894751.1   223100  ............................TC....  223067
PP770439.1   349135  ............................TC....  349102
PP035765.1   118497  ............................        118524
NC_086688.1  93086   ............................TC....  93053
MN946545.1   250268  ............................        250295
NC_057143.1  436419  ............................        436446
NC_057142.1  436410  ............................        436437
KF933043.1   333     ........                            340
```

Table S3. Estimation of detection limits of RT-LAMP assays for CTV and COX using RT-qPCR standard curves

| X-axis | Cq value (y) | SD | Derived X | Calculated copies per 2 uL | Calculated copies per 1 uL | Calculated copies per mg of tissue | |
|---|---|---|---|---|---|---|---|
| 6 | 18.5338 | 0.06013 | 6.033779 | 1080883.954 | 540441.9771 | 1080883.954 | |
| 5 | 22.11906 | 0.00845 | 4.991552 | 98073.64704 | 49036.82352 | 98073.64704 | |
| 4 | 25.36734 | 0.05397 | 4.047285 | 11150.25716 | 5575.12858 | 11150.25716 | |
| 3 | 28.91212 | 0.056 | 3.016826 | 1039.502604 | 519.7513018 | 1039.502604 | |
| 2 | 32.63547 | 0.09708 | 1.934456 | 85.9916724 | 42.9958362 | 85.9916724 | RT-LAMP detection limit |
| 1 | 36.19771 | 1.04151 | 0.898922 | 7.923581177 | 3.961790589 | 7.923581177 | |
| 0 | 37.99759 | 2.42965 | 0.375701 | 2.37520217 | 1.187601085 | 2.37520217 | |
| **CTV** | | | | | | | |
| | y = -3.44x + 39.29 | | | | | | |

| X-axis | Cq value (y) | SD | Derived X | Calculated copies per 2 uL | Calculated copies per 1 uL | Calculated copies per mg of tissue | |
|---|---|---|---|---|---|---|---|
| 6 | 15.02381 | 0.05403 | 5.858101 | 721274.4939 | 360637.247 | 721274.4939 | |
| 5 | 18.12891 | 0.02711 | 4.96583 | 92433.726 | 46216.863 | 92433.726 | |
| 4 | 21.50199 | 0.05848 | 3.996555 | 9920.980539 | 4960.490269 | 9920.980539 | |
| 3 | 24.91915 | 0.00777 | 3.014612 | 1034.217945 | 517.1089724 | 1034.217945 | |
| 2 | 28.54531 | 0.0107 | 1.972612 | 93.88842829 | 46.94421415 | 93.88842829 | |
| 1 | 31.90146 | 0.08333 | 1.008201 | 10.19063272 | 5.095316359 | 10.19063272 | RT-LAMP detection limit |
| 0 | 36.0439 | 1.00244 | -0.18216 | 0.6574229 | 0.32871145 | 0.6574229 | |
| **COX** | | | | | | | |
| | y = -3.48x + 35.41 | | | | | | |

- Derived X = (Cq value – intercept of the formula)/(slope of the formula)
- Calculated copies per 2 μL = 10^(Derived X); Per 2 μL because each reaction contains 2 μL nucleic acid template
- Calculated copies per 1 μL = (Calculated copies per 2 μL)/2
- Calculated copies per mg of tissue: The amounts of plant tissue and lysis buffer used for nucleic extraction are 250 mg and 750 μL. Then, 150 μL of supernatant out of 750 μL, equivalent to 1/5, was taken for benchtop extraction. Thus, the supernatant contains 50 mg (i.e. 250 mg/5) of plant tissue equivalently. All nucleic acids extracted from the supernatant were resuspended in 100 μL of elution buffer. Thus, every 2 μL of the elute contains nucleic acids equivalent to 1 mg of the plant tissue. Consequently, "per 2 μL" and "per mg of tissue" become interchangeable.

Table S4. Assessment of the RT-LAMP detection efficacy to various CTV isolates

| Category | Source rootstock | CTV isolates | GenBank accession no. | RT-qPCR | | RT-LAMP | |
|---|---|---|---|---|---|---|---|
| | | | | CTV | COX | CTV | COX |
| CTV-infected citrus species | Rough lemon | T500 | KC841779.1 | 3/3 | 3/3 | 3/3 | 3/3 |
| | Unknown | T500 | KC841779.1 | 3/3 | 3/3 | 3/3 | 3/3 |
| | Pineapple sweet orange | T517 | KC841785.1 | 3/3 | 3/3 | 3/3 | 3/3 |
| | Unknown | T520 | KC841786.1 | 3/3 | 3/3 | 3/3 | 3/3 |
| | Unknown | T521 | KC841811.1 | 3/3 | 3/3 | 3/3 | 3/3 |
| | Pineapple sweet orange | T525 | KC841821.1 | 3/3 | 3/3 | 3/3 | 3/3 |
| | Unknown | T529 | KC841789.1 | 3/3 | 3/3 | 3/3 | 3/3 |
| | Pineapple sweet orange | T534 | KC841792.1 | 3/3 | 3/3 | 3/3 | 3/3 |
| | M.V. sweet orange | SY554 | KC841799.1 | 3/3 | 3/3 | 3/3 | 3/3 |
| | Pineapple sweet orange | SY558 | KC841826.1 | 3/3 | 3/3 | 3/3 | 3/3 |
| | M.V. sweet orange | SY560 | KC841813.1 | 3/3 | 3/3 | 3/3 | 3/3 |
| | Pineapple sweet orange | SY568 | KC841814.1 | 3/3 | 3/3 | 3/3 | 3/3 |
| | M.V. sweet orange | SY579 | KC841796.1 | 3/3 | 3/3 | 3/3 | 3/3 |
| | Unknown | SY580 | KC841818.1 | 3/3 | 3/3 | 3/3 | 3/3 |

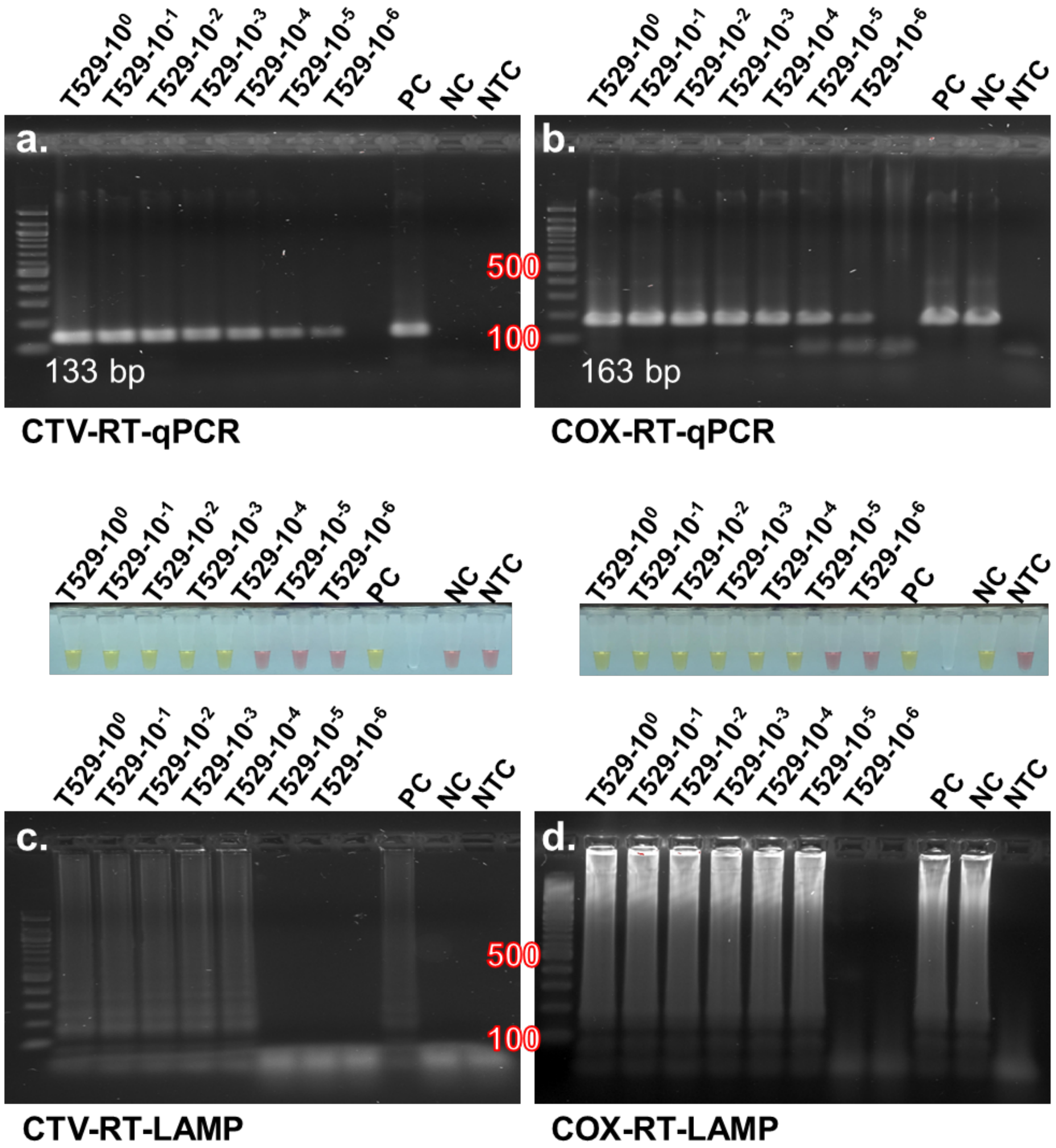


Fig. S1. Gel images of (a) RT-qPCR and (b) RT-LAMP assays using serial dilutions of a Citrus tristeza virus (CTV isolate). Gel images of (c) RT-qPCR and (d) RT-LAMP assays for detection of cytochrome oxidase (COX) reference gene. Inserted images above Fig. S1c. and S1d. show color changes of the reactions in Eppendorf tubes after RT-LAMP assays.

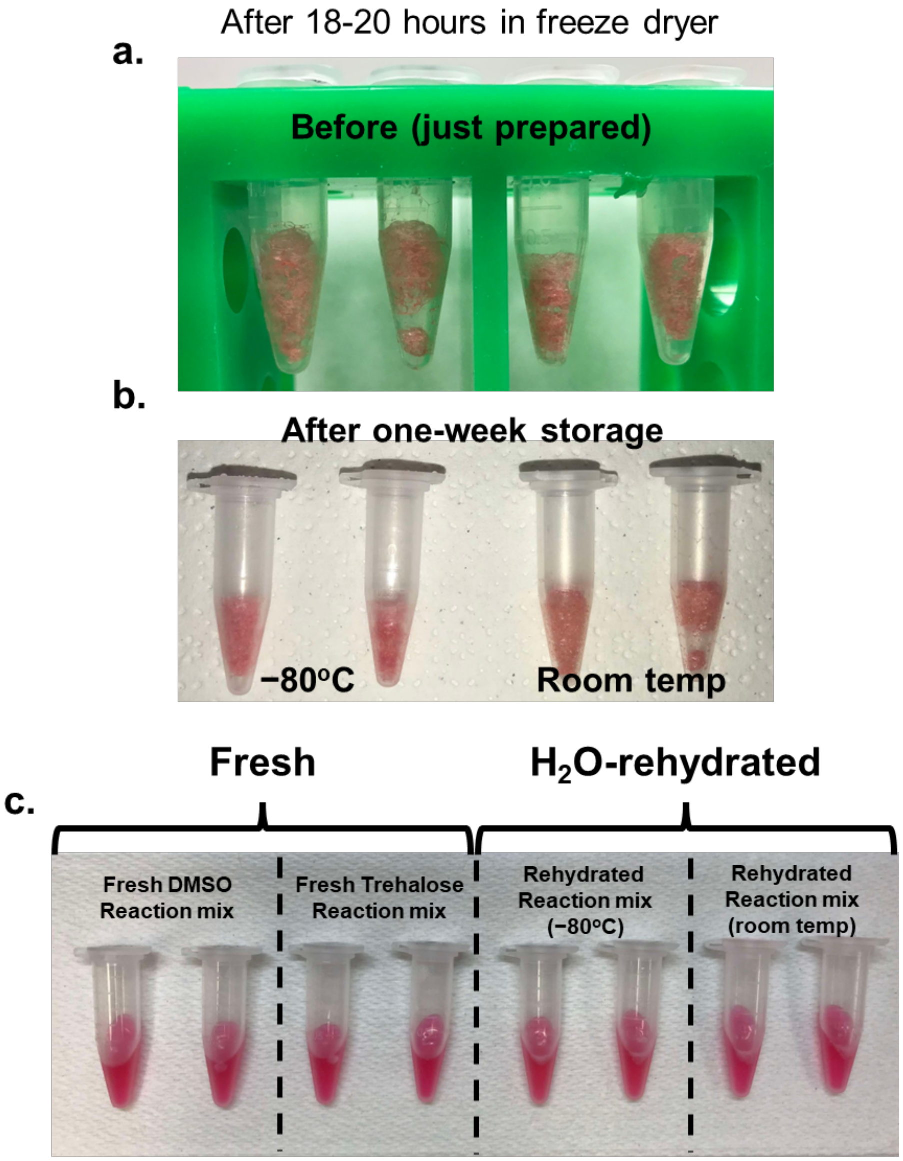


Fig. S2. Lyophilized reaction mix after 18-20-hour lyophilization process (a) freshly prepared and (b) after one week of storage. (c) The comparison between fresh controls and the rehydrated reaction mix, showing no distinguishable color difference.
